# Co-aggregation of pre-conditioned cell populations drives controlled lineage bias and formation of gut tube-like structures in human stem cell-based embryo models

**DOI:** 10.64898/2026.11.09.765272

**Authors:** Igor Meszka, Almira Ibragimova, Max Lycke, Natalia Smirnova, Sergey Ponomartsev, Rui Zhang, Tharvesh Moideen Liyakat Ali, Yuliia Boichuk, Brian K. Chung, Jonas Øgaard, Espen Melum, Nikolaj Gadegaard, Stefan Krauss

## Abstract

Human stem cell-based embryo models (hSCBEMs) provide a powerful platform for investigating post-implantation human embryo development but are frequently hindered by variability in growth and morphogenesis as well as incomplete lineage representations. Here, we present a modular hSCBEM aggregation approach based on co-aggregation of differentially pre-conditioned, human pluripotent stem cell derived cells pools. Using this strategy, we generated hSCBEM with consistent morphotypes that targets robust morphogenesis of gut tube-like structures in combination with mesodermal lineages. Our findings demonstrate that co-aggregating defined cellular heterogeneity represents a versatile approach to specifically guide self-organization in hSCBEMs while improving reproducibility and developmental fidelity. The approach provides a robust platform for supervised morphogenesis in hSCBEMs.

## Introduction

Understanding early human development remains constrained by limited access to post-implantation human embryos and the associated ethical considerations (Rossant and Tam, 2021; Rugg-Gunn et al., 2023; Rivron et al., 2023; Bentzen et al., 2026). Stem cell-based embryo models (SCBEMs) have therefore emerged as a powerful experimental platform for investigating peri-and post-implantation development and pathogenic concepts (Pera et al., 2015; Rivron et al., 2018; Moris et al., 2020; Moris et al., 2026; Martinez-Arias et al., 2026). Generated from pluripotent stem cells, these systems recapitulate key aspects of mammalian embryogenesis while providing experimental accessibility, scalability, and opportunities for mechanistic perturbation (Moris et al., 2026; Martinez-Arias et al., 2026).

3D gastruloids represent one of the widely used classes of SCBEMs, recapitulating key developmental processes including symmetry breaking, axial organization, and specification of the three germ layers (van den Brink et al., 2014; Beccari et al., 2018; Moris et al., 2020; Dingare and Steventon, 2023; Martinez Arias et al., 2022; Turner and Martinez-Arias, 2024; Liu and Wu, 2025). Human gastruloid technology relies on the aggregation of a defined number of human pluripotent stem cells (hPSCs) with a transient pulse of the GSK3 inhibitor CHIR99021 during 2D culture pre-treatment and in the first 24 hours after aggregation. This generates structures that recapitulate features of Carnegie stage 9 embryos (approx. 19-21 days post fertilization) (Moris et al., 2020). Additional engineering of the extracellular environment and/or signaling landscape enables complex post-gastrulation developmental programs such as neural patterning (Olmsted and Paluh, 2021), cardiac induction (Olmsted and Paluh, 2022), and somitogenesis (Sanaki-Matsumiya et al., 2022; Miao et al., 2023; Yamanaka et al., 2022; Hamazaki et al., 2024).

The reliance on uniform WNT activation carries a consequence that defines the current limits of the approach: the entire aggregate is driven toward posterior identity, and anterior structures such as the foregut and the anterior neural tube are not obtained (Liu et al., 2025). A second, less examined consequence concerns the endoderm. Representation of definitive endoderm in both mouse (Veenvliet et al., 2020; Rossi et al., 2021; Hashmi et al., 2022), and human gastruloids (Hamazaki et al., 2024), has remained limited and inconsistent, and a continuous, regionalized gut tube has not been described as a reproducible feature of a human model. This matters because gut endoderm is the substrate for the organogenesis of the respiratory and digestive systems, and because its anteroposterior regionalization depends on signals from the adjacent mesoderm — precisely the kind of tissue–tissue interaction that embryo models are best placed to interrogate.

The lineage composition and robustness of human gastruloid generation depends on multiple experimental parameters, including the specific hPSC line, 2D culture conditions such as cell confluency, colony size, oxygen pressure, metabolic state and passaging (Moris et al., 2020), as well as perturbations of signaling pathways such as WNT and NODAL during the pre-treatment and aggregation stages (Neupane et al., 2025; Azami et al., 2025; Das et al., 2026).

Recent studies in mouse gastruloids suggest that intrinsic heterogeneity within the initial pluripotent stem cell culture might be a further key determinant of gastruloid formation, influencing symmetry breaking, cell fate specification, and inter-experimental variability (Farag et al., 2024; Anlas et al., 2025; Ayyappan et al., 2026; Blotenburg et al., 2025). A heterogeneity based differential responsiveness to CHIR99021-mediated WNT activation has been proposed to facilitate symmetry breaking through cell sorting and self-organization, enabling anterior-posterior axis formation in the absence of externally imposed morphogen gradients or spatial cues (McNamara et al., 2024). While controlled culture conditions can modulate the extent of this intrinsic heterogeneity, lineage potential and subsequent composition of gastruloids, (Cermola et al., 2021; Anlas et al., 2025), it remains difficult to achieve controlled morphotypes when starting from a single PSC founder population.

Here, we developed an alternative strategy for human gastruloid aggregation based on the introduction of multiple defined cellular pools prior to aggregation. We generated a series of pre-conditioned human embryonic stem cell (hESC) populations by differentially modulating WNT and TGFβ signaling, thereby creating cell states with distinct differentiation biases and lineage competencies. We hypothesized that combining these populations during aggregation would promote the responses required for symmetry breaking by simultaneously engaging both the anterior and posterior primitive streak regulatory modules recently described in mouse gastruloids (Dias et al., 2025; Martinez-Arias et al., 2025). The hSCBEM composed of three pools of differentially pre-treated cells, which we termed aggregoids, exhibited robust formation of morphologically stable structures and reproducible endoderm morphogenesis. Together, these findings establish engineering of controlled cellular heterogeneity as a strategy to direct developmental complexity while improving reproducibility, and morphogenetic fidelity of human gastruloids.

## Results

### Pre-treatment creates fate-biased cell populations

To generate controlled heterogeneity in the starting material for hSCBEMs, we tested and characterized four pre-treatment conditions, each producing a cell population with distinct expression profiles and developmental biases prior to aggregation. Untreated (UT) cells represented hESCs (RUES2-GLR) (Martyn et al., 2018) in a defined pluripotency maintenance medium E8 (A1517001, Life Technologies) containing insulin, FGF2 and TGFβ1 (Chen et al., 2011). In the CHIR pre-treatment condition, WNT signaling was activated by supplementing E8 medium with CHIR99021. To reduce cell cycle-dependent variability in the CHIR99021 response (Laco et al., 2018), hESCs were synchronized by transient nocodazole treatment prior to CHIR99021 treatment (Figure 1, Methods). To generate a population with an enhanced mesendodermal bias, cells were exposed to the RACL pre-treatment condition (Linneberg-Agerholm et al., 2019, Methods), which activate WNT, NODAL, and LIF signaling in the absence of insulin (Methods). In the TNKSi pre-treatment condition, hESCs in E8 were exposed to the Tankyrase1/2 inhibitor OM-153 (Waaler et al., 2020) to target endogenous WNT and Hippo signaling.

Cell colonies in the CHIR pre-treatment first underwent compaction after Nocodazole exposition, an event that was then reversed upon the addition of CHIR99021 – colonies then started to have more rounded edges and obtained a dome-like shape. In RACL pre-treatment, the typical smooth-edged, compact morphology of RUES2-GLR colonies, composed of small cells with a high nucleus-to-cytoplasm ratio, shifted to colonies with irregular, spiky borders consisting of flatter, more polygonal cells and reduced cell compaction (Figure 1A). In contrast, TNKSi pre-treatment resulted in more compact hESC colonies with enhanced cell adhesion and higher resistance to enzymatic detachment during passaging (Figure 1A).

To assess the impact of pre-treatment conditions while tracking the heterogeneity of the pretreated cells, we performed single-cell RNA sequencing (scRNAseq) of the 2D cultures before aggregation. The analysis revealed substantial overlap between the UT and TNKSi populations in the UMAP space, whereas the CHIR and RACL populations were substantially segregated (Figure 1B, C). Across all samples, we identified seven distinct clusters by unsupervised, graph-based clustering of the sequenced cells (Figure 1B). Cell populations corresponding to UT and TNKSi were annotated with clusters 1, 4 and 7, whereas the RACL sample was represented by clusters 2 and 5; the CHIR pre-treated population consisted of clusters 3 and 6 (Figure 1C).

Both CHIR-specific clusters 3 and 6 expressed canonical WNT target genes, including *DKK1, SP5, TCF7,* and *LEF1*, as well as WNT-dependent posterior markers such as *CDX1* and *TBXT*, consistent with a primitive streak-like state. However, the two clusters differed in the expression of *AXIN2* and *CDX2*, which were upregulated in cluster 6 but not cluster 3, suggesting heterogeneity in the cellular response to the CHIR99021 treatment (Figure 1D). RACL-specific clusters 2 and 5 exhibited an early endodermal-like transcriptional signature characterized by upregulated expression of NODAL-associated genes, including *EOMES, LHX1, FOXA2, GSC,* and *LEFTY1*, especially pronounced in cluster 2 (Figure 1D). Both CHIR and RACL pre-treated populations contained a substantial TBXT-positive cell fraction, however, only RACL cells showed a *SOX17*::mCherry signal in a salt-and-pepper manner, while in the CHIR condition TBXT extensively co-localized with *SOX2::*mCitrine (Figure 1E).

TNKSi and UT pools were composed predominantly of *SOX2*::mCitrine positive cells with a pluripotency-associated transcriptional signature (Figure 1D, E). Cluster 4, which was prominent in TNKSi-treated cells, exhibited elevated expression of Hippo/YAP-associated transcriptional regulators, including *TEAD2, TEAD3, TEAD4,* and *YAP1*. In addition, cluster 7, displayed increased expression of HIPPO pathway regulatory genes, including *LATS1, LATS2,* and *YAP1*, suggesting that TNKSi pre-treatment is associated with altered regulation of the Hippo-YAP pathway (Wang et al., 2015).

UT, TNKSi, and CHIR cells retained predominantly CDH1-positive epithelial phenotype (Figure 1E) and expressed epithelial markers including *CDH1, EPCAM, CLDN6,* and *TJP1* (Figure 1D). In contrast, the RACL-associated clusters exhibited upregulation of markers for epithelial-to-mesenchymal transition (EMT) *CDH2, VIM, FN1, ITGA5,* and *MMP14*, together with a reduced expression of epithelial markers consistent with the initiation of EMT during anterior primitive streak specification (Williams et al., 2011, Sheng et al., 2021) (Figure 1D).

Together, these results demonstrate that the four pre-treatment conditions generated transcriptionally distinct cell populations resembling pluripotent, posterior primitive streak-like, and early endodermal states respectively and providing a tunable starting material for hSCBEM generation.

**Figure 1.**
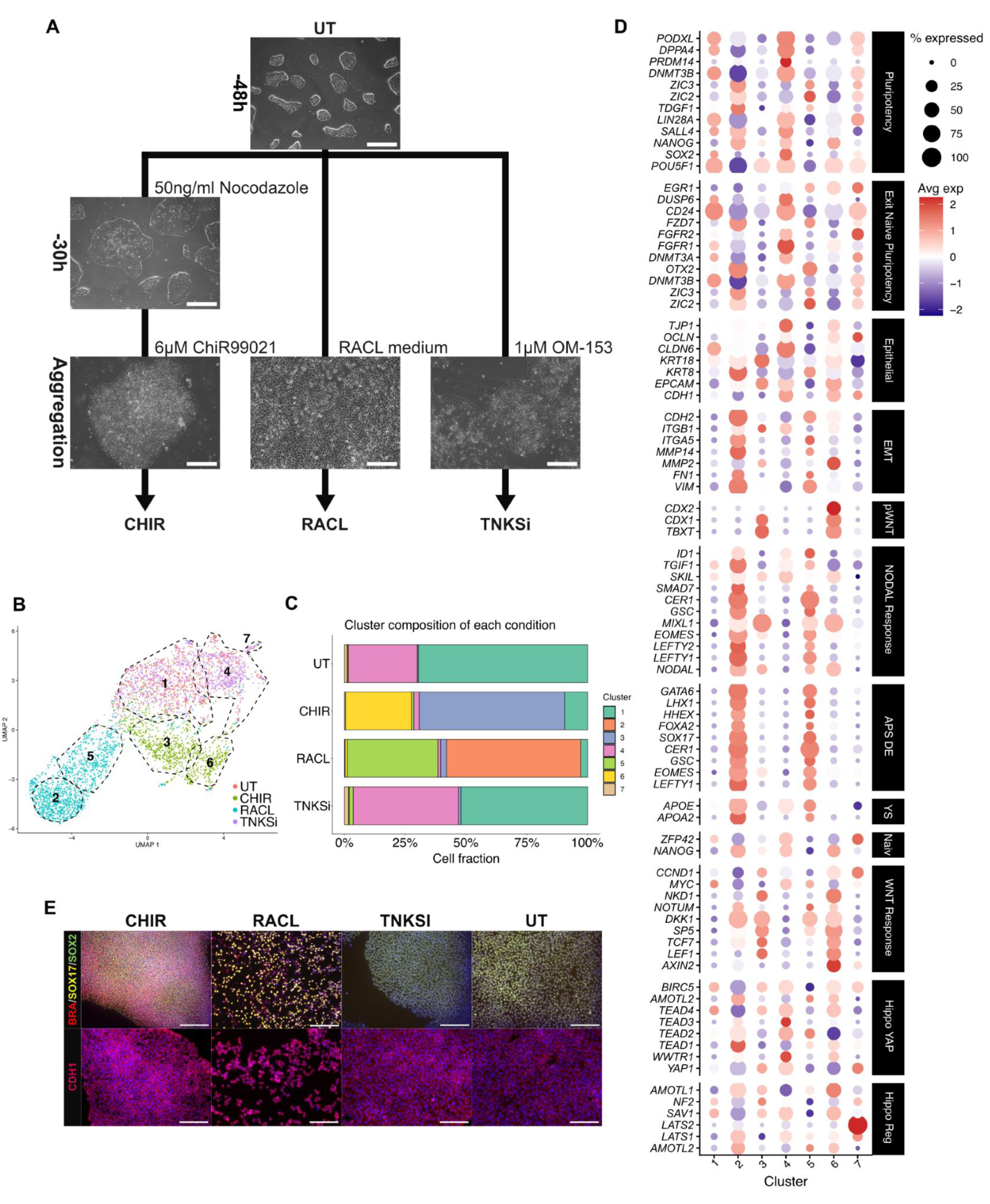
Pre-treatment creates fate-biased cell populations. **A)** Phase contrast images of RUES2-GLR hESC 2D culture under pluripotency maintaining conditions (UT) and upon pre-treatment with CHIR, RACL and TNKSi conditions. Scale bar: 550 µm. **B)** UMAP visualization based on scRNA-seq analysis of the four 2D pre-treated samples, showing distinct pools of cells color-coded by condition. Outlined regions mark distinct annotated cell populations identified by unsupervised clustering. **C)** Stacked bar plot showing the proportions of cell clusters within each 2D cell pool. The pre-treatment color-coding matches Figure 1B. **D)** Dot-plot showing averaged expression and percentage of cells expressing selected marker genes across identified clusters in “UT”, “CHIR”, “RACL” and “TNKSi” pools. **E)** Representative immunofluorescent images of BRA (also known as TBXT), *SOX2*::mCitrine and *SOX17*::mCherry, and CDH1 distribution in pretreated 2D cultures. Scale bar: 550 µm

### Co-aggregation strategies lead to the emergence of reproducible hSCBEM morphotypes

Next, we assessed the potential of the pre-treated cell populations to form hSCBEMs, either alone or in defined combinations. The pre-treated cell pools and combinations of them were aggregated at 600 cells per well, in total, and cultured in E6 medium supplemented with 1 μM CHIR99021 and 2 μM SB431542, followed by culture in E6 medium for five days (Figure 2A, Methods).

Among all tested conditions, only CHIR cells consistently formed elongated aggregates resembling classical gastruloids (Moris et al., 2020) (Supplementary Figure 1A). Both RACL and TNKSi cell pools alone did not exhibit symmetry breaking or axial elongation (Supplementary Figure 1A). However, some of the double, triple or quadruple mixes of CHIR cells with UT, TNKSi and RACL cells (Supplementary Figure 1 B) led to elongated structures (Supplementary Figure 1C) to which we refer as aggregoids. In the further work we focused on the double (two lineage) 1:1 mixture of CHIR-, RACL-pretreated cells and the triple (three lineage) mix 1:1:1 mixture of CHIR-, RACL-, and TNKSi-pretreated cells (Figure 2A) based on its ability to form reproducible morphotypes with posteriorly localized TBXT and *SOX2::*mCitrine expression and a distinct *SOX17*::mCherry-positive continuous structure (Figure 2B, C, Supplementary Figure 1D, E).

Successful symmetry breaking and axial elongation were observed in 73% of the CHIR, TNKSi and RACL triple-mix aggregoids (N=960; n=10 independent experiments); 16% either failed to elongate or exhibited abnormal morphology (classified as “Other”; Supplementary Figure 2C), and 7% were excluded from the analysis for different reasons (classified as “Ignore”; Supplementary Figure 2C). Almost immediately after seeding, cells formed compact aggregates with distinctive *SOX17*::mCherry and *SOX2*::mCitrine domains (Figure 2B). Of interest, even when initially dispersed, coherent *SOX17*::mCherry positive populations were able to migrate, merge and form a single continuous domain by 72h post-aggregation.

Based on the expression of*SOX17*::mCherry, we identified two major distinct morphotypes in the triple-mix aggregoids: the “Inner” morphotype (61%) contained a lumenized *SOX17*::mCherry-positive tubular structure in the center of the hSCBEM, whereas the “Edge” morphotype (39%) displayed the *SOX17*::mCherry-positive domain at the periphery of the structure (Figure 2B, C, D). The two morphotypes were distinguishable as early as 48h after aggregation. In the “Inner” morphotype, the *SOX17*::mCherry positive domain was centrally positioned within the aggregoid and progressively extended toward the posterior end during elongation (Figure 2B, F upper row). In contrast, the “Edge” morphotype retained a peripheral *SOX17*::mCherry domain extending along the length of the structure from 48 to 120 h (Figure 2B, F lower row).

Morphometric analysis of the hSCBEM culture from day 2 to day 5 revealed that both triple-mix morphotypes exhibited a continuous increase in area and midline length, with progressively increasing inter-aggregoid variability over time (Figure 2E). The analysis of midline length across independent experimental plates and time points revealed that the “Inner” morphotype followed a remarkably consistent growth trajectory, whereas the “Edge” morphotype exhibited substantially greater variability and overall irregular growth dynamics (Supplementary Figure 2A, B). The occurrence of the “Inner” and “Edge” morphotypes varied depending on the position within the 96-well plate, with the “Inner” morphotype prevailing in the central and inner-boundary wells (50% vs. 26% and 52% vs. 22%, respectively), whereas both morphotypes occurred at similar frequencies in outer-boundary wells (39% each) (Supplementary Figure 2D, E), suggesting micro-environmental differences in the 96-well plate that influence morphotype specification.

In summary, we show that aggregating hSCBEMs from specific, pre-differentiated cell pools leads to controllable and reproducible differentiation. We observed stable growth and patterning, and based on *SOX17*::mCherry expression, classified two distinct morphotypes.

**Figure 2.**
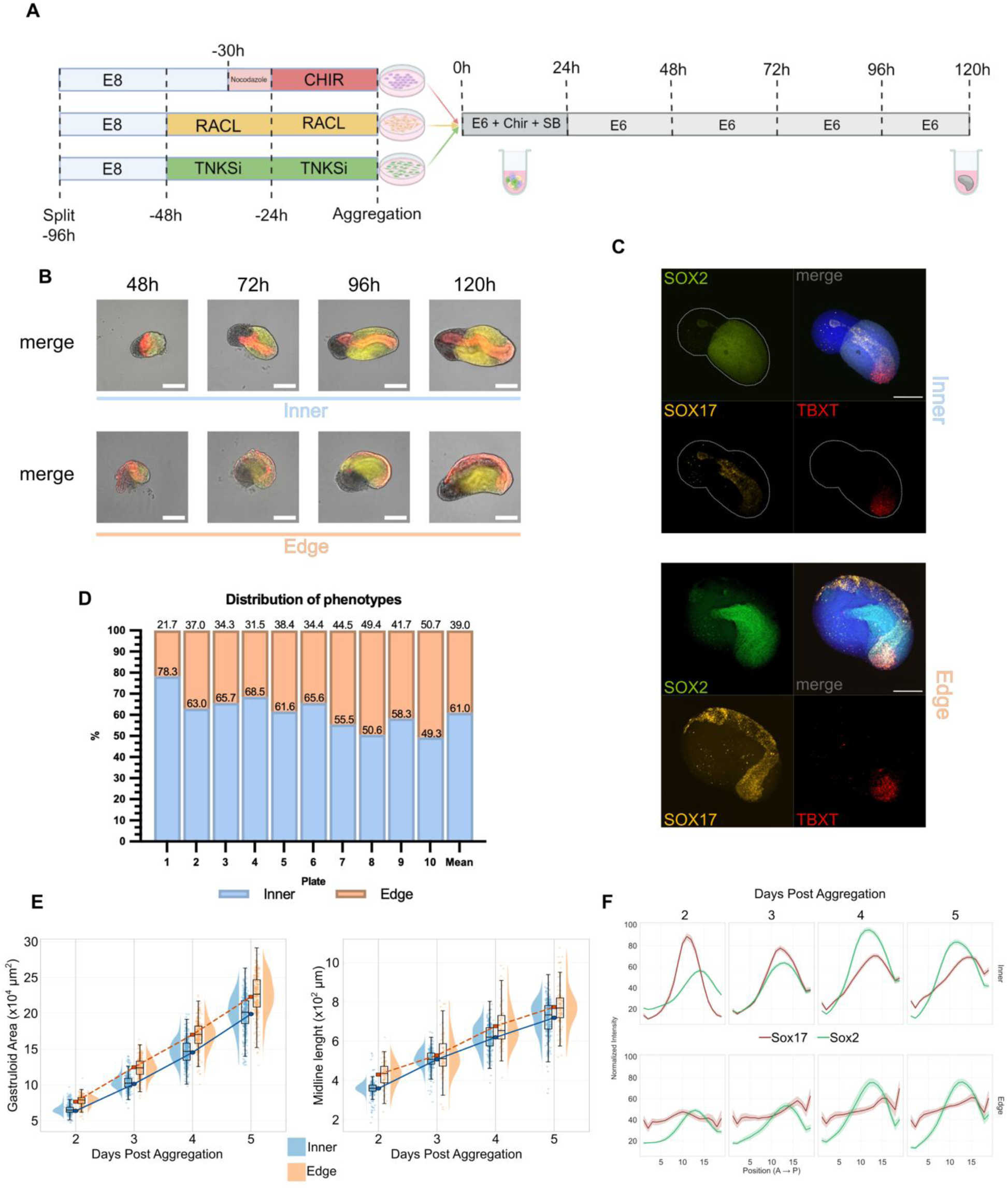
Co-aggregation strategies lead to emergence of reproducible hSCBEM morphotypes. **A)** Schematic overview of the experimental workflow for generating hSCBEMs by co-aggregation of pre-differentiated cell populations, followed by culture in CHIR99021 and SB431542-containing medium. **B)** Representative images of hSCBEMs displaying “Inner” and “Edge” morphotypes across a 5-day cell culture, with overlayed signal from fluorescent reporters *SOX17*::mCherry and *SOX2*::mCitrine. Scale bar: 200 μm. **C)** Immunofluorescent images demonstrating germ layer organization of triple-mix aggregoids at 120h. Scale bar: 200 μm. **D)** Quantification of hSCBEM morphotypes across independent experiments, showing relative frequencies of “Inner” and “Edge” morphotypes across successful hSCBEMs. **E)** Quantitative morphometric analysis of hSCBEM development over time. Left: aggregate area progression. Right: axial elongation measured by midline length. Data are stratified by phenotypes (“Inner” vs “Edge”). **F)** Normalized *SOX2*::mCitrine and *SOX17*::mCherry fluorescent intensity profile along midline by day and morphotype. The X-axis represents the relative slice position from anterior to posterior.

### Single-cell transcriptomics reveals tunable cell type diversification in hSCBEMs

To assess cell type composition of the resulting hSCBEMs, we performed single-cell RNA sequencing (scRNA-seq) of double and triple-mix (representing both “Inner” and “Edge” morphotypes) aggregoids across four time points (48, 72, 96 and 120 h). In summary, data from 13076 cells in triple-mix “Inner” morphotype, 11640 in triple-mix “Edge” morphotype aggregoids and 15686 in double-mix aggregoids were analyzed across four time points.

At 48h (day 2 of culture) both “Inner” and “Edge” triple-mix aggregoids consisted of *NKX1-2, CDX2, TBXT* and *SOX2*-expressing population identified as neuromesodermal progenitors (NMPs), a major fraction of *MSGN1* and *TBX6*-expressing cells classified as pre-somitic mesoderm (PSM), a *RIPPLY2*/*MESP2*-expressing population resembling the determination front, *PAX8*, *WT1* and *OSR1*-expressing cells corresponding to intermediate mesoderm (IM) and *SOX17/EPCAM-*expressing cells annotated as gut endoderm (Figure 3A, C-F). At 72h (day 3), a *HAND1* and *PRRX1*-expressing population resembling lateral plate mesoderm (LPM) appeared along with a minor *TBX1*-positive cluster annotated as anterior pharyngeal mesoderm (APM). Within paraxial mesoderm population, two clusters of somitic mesoderm (SM) emerged, probably resembling earlier (*MEOX1-*high*, MEOX2-*low*, PAX3-*low) and more advanced (*MEOX1-*high*, MEOX2-*high*, PAX3-*high) somite-like populations (Figure 3A, C-F). At this stage, gut endoderm started differentiation towards *CDX2* and *APELA*-expressing cells indicating hindgut development (for both “Inner” and “Edge” morphotypes) and *FOXA1* and *IGFBP6-*expressing cells indicating midgut development (in “Edge” morphotype only) (Figure 3A, C, D). At 96-120 h, we observed a trend towards expansion of SM clusters at the expense of a shrinking PSM population, accompanied by an expansion of midgut and hindgut populations and a corresponding decrease of gut endoderm, likely reflecting progressive differentiation and maturation within the hSCBEMs (Figure 3E). Notably, the NMP population and a small cluster of *CD34, KDR, CDH5* and *SOX17*::mCherry-positive endothelial cells persisted throughout the culture period and remained well represented at comparable proportions across all time points from 48 to 120 h (Figure 3 A-E). The neuroectodermal lineage was represented by a minor *SOX2/SOX10*-positive cluster annotated as neural crest (NC), suggesting a strong mesodermal bias of NMPs (Figure 3A-E).

Immunostaining of triple-mix aggregoids revealed NMPs, identified as SOX2/TBXT double-positive cells, at the posterior CDX2-positive pole (Supplementary 3A-B). By 120 h, the anterior region of triple aggregoids contained somitic mesoderm (MEOX1-positive) (see Supplementary 3F), intermediate mesoderm (WT1-positive), and lateral plate mesoderm (FOXF1-positive)-like cell types (Supplementary 3D-E). EPCAM/SOX17 double-positive cells formed a structure resembling a gut tube (Supplementary 3C-D). Notably, most *SOX17*::mCherry-positive cells within this structure also expressed *SOX2::*mCitrine, suggesting an immature or transitional endodermal state (Supplementary 3D). A small number of *SOX17*::mCherry-positive but EPCAM-negative cells at the anterior tip of the structure may represent endothelial-like cells (Supplementary 3C).

At 48 h, double-mix aggregoids exhibited a cell-type composition similar to that of triple-mix aggregoids, comprising gut endoderm, anterior/segmenting PSM, intermediate mesoderm (IM), PSM, and a smaller fraction of somitic mesoderm (SM) cell types (Figure 3B, E, F). However, in contrast to triple-mix aggregoids, double-mix aggregoids contained almost no NMPs. At 72 h, the proportion of PSM cells had already substantially decreased and had almost disappeared by 96 h, coinciding with the expansion of SM cell types (Figure 3E, F). Notably, at 120 h, a substantial *PAX1/PAX9/SOX9*-expressing population resembling sclerotome emerged, suggesting a ventralization and further differentiation of somitic mesoderm (SM) (Figure 3A-B, E). Unlike triple-mix aggregoids, gut endoderm in double-mix aggregoids predominantly differentiated towards a midgut rather than hindgut identity, indicating a lower degree of posteriorization (Figure 3B, E).

Together, these results demonstrate that a modular engineering approach based on the co-aggregation of pre-treated cell populations enables the generation of hSCBEMs with tunable cell-type compositions and developmental trajectories.

**Figure 3.**
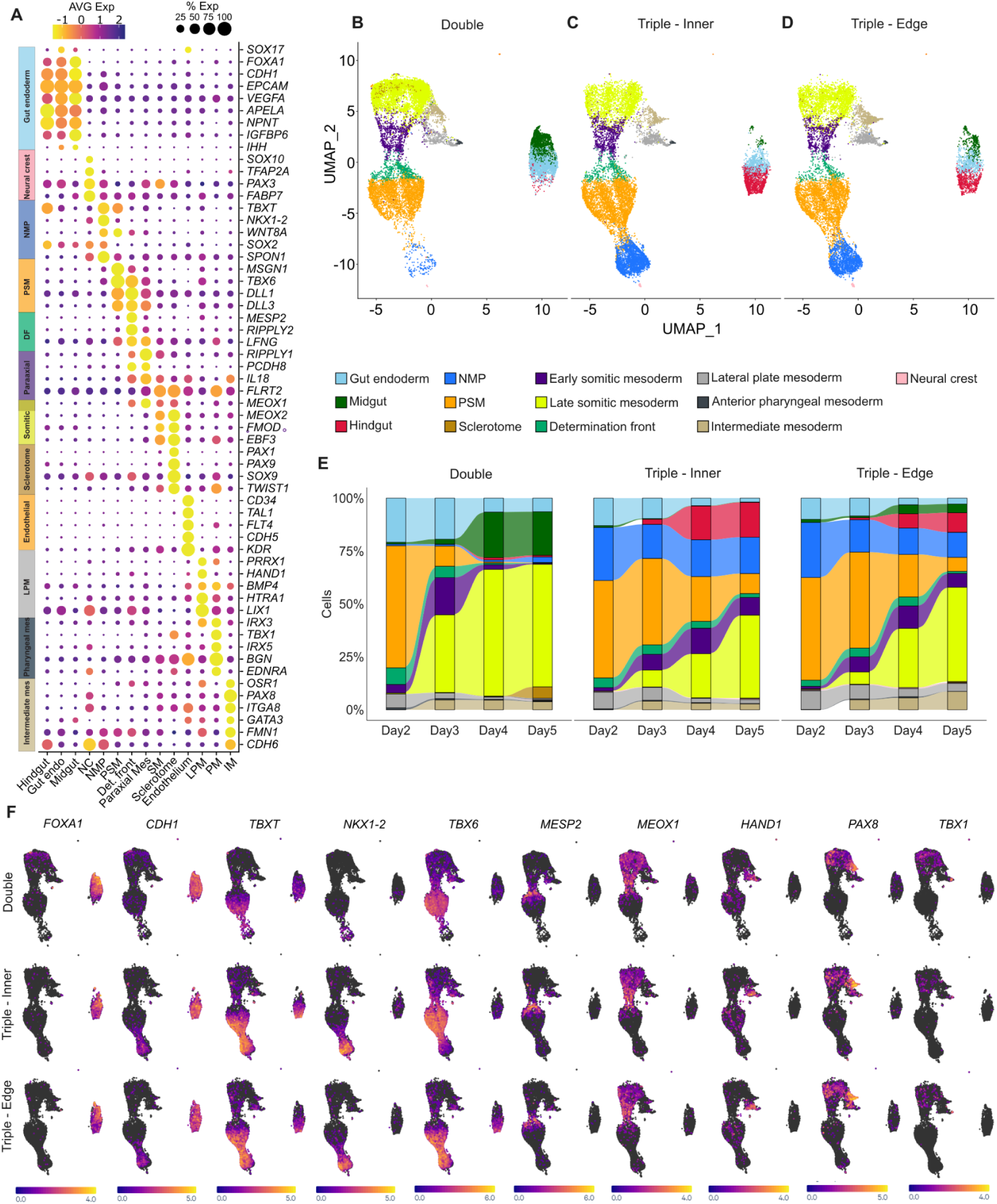
Analysis of cell type diversity in aggregoids by single-cell transcriptomics. **A)** Dotplot representation showing the expression of selected markers across various cell types annotated in all hSCBEM aggregoids. **B-D)** Uniform manifold approximation and projection (UMAP) of aggregated dataset of all time points divided by phenotypes. Double 48h = 5168, 72h = 3848, 96h = 4356, 120h = 2314 cells. Inner 48h = 3626, 72h = 3110, 96h = 2183, 120h = 4157 cells. Edge 48h = 2406, 72h = 2695, 96h = 2937, 120h = 3602 cells. **E)** Stacked area plots representing changes in cell composition in various phenotypes color-coded by clusters identity. **F)** UMAP feature projection of selected gene expressions in aggregated dataset of all time points divided by phenotypes.

### Comparative analysis of hSCBEM cell composition reveals improved gut-tube formation and gut-endoderm regionalization

To analyze the lineage composition of the double and triple-mix aggregoids, including “Inner” versus “Edge” phenotypes, we compared and integrated our scRNA-seq datasets (120h endpoint; n = 2314, 4157 and 3602 cells, for “Double”, “Inner” and “Edge”, respectively) with a reference human embryo dataset at post-conceptional week 3 (PCW3) corresponding to Carnegie stage 10 (CS10) (*n* = 10,444 cells; Zeng et al., 2023) (Figure 4A). Canonical human gastruloids (96 h) and a retinoic acid (RA)-induced somitic model (96–120 h) (Hamazaki et al., 2024), together with our aggregoid datasets, were projected onto this reference by anchor-based label transfer and UMAP projection (MapQuery, Seurat v5), with transferred labels subsequently verified by marker-based gating (Figure 4B–F; Methods). Transcriptional correspondence between aggregoid and embryo cell types was quantified separately by correlation of pseudobulk expression profiles (Supplementary Figure 4; Methods). All datasets were re-annotated under a common lineage schema to enable direct cross-dataset comparison (see Methods). Overall, our hSCBEMs aligned well with the PCW3 human embryo reference, indicating that they recapitulate sets of major developmental trajectories of early human embryogenesis (Figure 4D-F). Furthermore, correlation analysis demonstrated a high degree of similarity between most of the identified cell populations and their corresponding counterparts in the human embryo reference (Supplementary 4A). Nevertheless, compared with the PCW3 human embryo reference, both double and triple-mix aggregoids exhibited limited neuroectodermal trajectories, only a sparse population of notochord-like cells, underrepresented cardiac mesoderm, and an absence of hemogenic cell types. Double and triple-mix aggregoids exhibited an overall cellular organization comparable to that of previously published canonical human gastruloids at 96 h (Hamazaki et al., 2024) while containing an intermediate mesoderm (IM) population as well as populations of midgut, and sclerotome cells found predominantly in double-mix aggregoids (Figure 4G). In contrast to the 96–120 h RA-induced somitogenic gastruloids (RA gastruloids) (Hamazaki et al., 2024), double and triple-mix aggregoids retained a substantial NMP population but lacked neural tube and cardiomyocytes (Figure 4C-G). The most prominent feature distinguishing both double and triple-mix aggregoids from canonical and RA gastruloids was the robust formation of extensive endodermal populations and the ability to generate a sclerotome-like population (Figure 4D-G).

Overall, we show that the modular engineering approach enables the generation of hSCBEMs that recapitulate key developmental features of the PCW3 human embryo and exhibit developmental potential comparable to existing hSCBEMs, while providing distinct advantages for modeling endoderm development.

**Figure 4.**
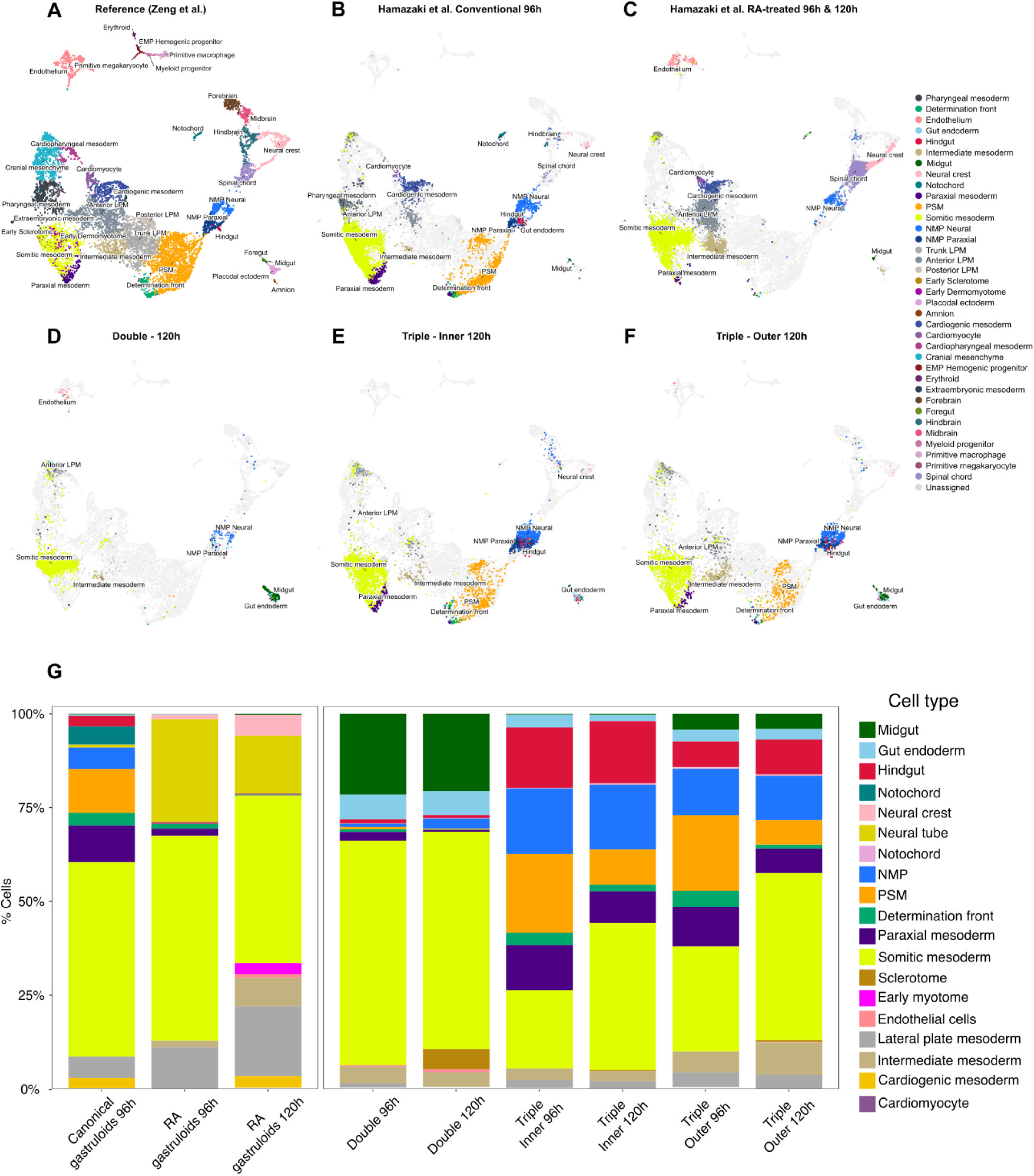
Comparative analysis of hSCBEM cell composition reveals enhanced endoderm development. **A)** UMAP of the PCW3 (CS10) human embryo reference (*n* = 10,444 cells; Zeng et al., 2023), coloured by curated reference annotation. **B, C)** Cells from canonical human gastruloids at 96 h (B) and RA gastruloids at 96–120 h (C) (Hamazaki et al., 2024), projected into the reference embedding by anchor-based label transfer and UMAP projection (MapQuery, Seurat v5) after marker-based gating (Methods). **D–F**) Cells from double-mix (D), triple-mix “Inner” (E) and triple-mix “Edge” (F) aggregoids at 120 h, projected and coloured as in B and C. In all query panels, gut regional identities (gut endoderm, midgut, hindgut) are taken from each dataset’s own annotation rather than transferred, because the reference contains too few gut cells for reliable regional label transfer (Methods). **G**) Stacked bar plot of cell-type composition across systems: canonical gastruloids (96 h), RA gastruloids (96 and 120 h) and aggregoids (96 and 120 h). Cell types that did not reach 0.1% of cells in any of the systems shown were excluded, as were cells flagged as artefacts.

### Discussion

Gastruloids recapitulate key morphological and molecular events of early embryogenesis, including the establishment of the three germ layers and the primary body axes (Beccari et al. 2018; van den Brink et al. 2014; Moris et al. 2020). In mouse gastruloids, which are normally made of naive embryonic pluripotent stem cells (Cermola et al. 2021), symmetry breaking and lineage specification processes are triggered by a single 24 h-long pulse of GSK3 inhibitor CHIR99021 given at 48 h after aggregation, when cells in aggregate reach epiblast-like state (van den Brink et al. 2014; Dias et al. 2025). In contrast, published human gastruloid protocols (Moris et al. 2020; Hamazaki et al., 2024; Azami et al. 2026) use primed human pluripotent stem cells (hPSC) as the starting material and require a 24h pre-treatment of hPSC 2D culture followed by CHIR99021 stimulation during the first 24h after aggregation. The pre-aggregation conditions of 2D culture such as cell density and colony size are shown to be critical for cell composition and elongation metrics of resulting gastruloids (Moris et al. 2020; Azami et al. 2026). Furthermore, studies on mouse gastruloids suggest that symmetry breaking depends on pre-existing signaling heterogeneity among cells within the aggregate, leading to differential response to CHIR stimulation (McNamara et al. 2024). Therefore, we sought to test whether combining cell populations with distinct biases, introduced at the pre-treatment step, could provide a reproducible and tunable strategy for human gastruloid formation and controlled modulation of their lineage composition.

In this proof of principle study, hESC populations were separately preconditioned with CHIR, RACL, or TNKSi inhibitor respectively, and then co-aggregated to form hSCBEM (aggregoids) exhibiting key features of human gastruloids. The resulting structures demonstrated anteroposterior axis establishment and axial elongation, as well as paraxial mesoderm formation and the emergence of two morphotypes of a gut tube-like structure (“Inner” and “Edge”). The comparison of scRNAseq profiles from triple-mix (CHIR, RACL, TNKSi) and double-mix (CHIR, RACL) aggregoids with a human Carnegie stage 10 (CS10) human embryo reference dataset (Zeng et al. 2023) showed substantial transcriptional similarity between clusters in SCBEMs and cell populations in three weeks post-conception embryo.

Similar to previously reported human gastruloids (Moris et al. 2020; Hamazaki et al. 2024; Azami et al. 2026), both double and triple-mix aggregoids exhibited a predominantly post-occipital body plan and lacked anterior neural structures, that, hence, reduced ethical concerns associated with the formation of more complete SCBEM, while remaining consistent with the principles of proportionate, model-specific ethical oversight and embedded ethics (Bentzen et al. 2026). Notably, our aggregoids exhibited a substantial enrichment of endodermal cell populations (gut endoderm, hindgut, midgut) compared with the reported models (Moris et al. 2020; Hamazaki et al. 2024). This effect was likely attributable to the shown pro-endodermal bias of RACL pre-conditioned cells.

Another distinguishing feature of triple-mix, but not double-mix aggregoids, was the persistence of neuromesodermal progenitors (NMPs) and significant amounts of early paraxial mesoderm cell types (presomitic mesoderm, early somitic mesoderm, determination front) at 120h post-aggregation. Although NMPs were scarcely represented in double-mix aggregoids already at 48h post-aggregation, we hypothesize that these progenitors were not entirely absent but rather emerged transiently at early stages and underwent rapid differentiation, leaving their downstream derivatives such as somitic mesoderm. This possibility was supported by the *TBXT/SOX2* double-positive profile of CHIR-preconditioned cells and the presence of putative NMP-derived populations at 48h, suggesting that the NMPs may have been exhausted before the first analyzed time point. The persistence of NMPs in triple-mix aggregoids suggested that the addition of TNKSi-pre-treated cells might have contributed to the prolonged maintenance of the NMP pool. Triple (but not double-mix) aggregoids also exhibited posterior patterning of the gut endoderm, especially the “Inner” morphotype. Although further experimental validation is required, our gene expression data, together with previous studies, suggested that gut posteriorization might have been influenced by proximity to *WNT3A*- and *WNT8A*-expressing NMPs (Schnirman et al. 2023) and presomitic mesoderm. This could explain the particularly pronounced endoderm posteriorization in the “Inner” morphotype, where the gut tube was surrounded by these populations, whereas in the “Edge” morphotype this exposure could be limited, giving a more balanced midgut/hindgut ratio. Notably, somitic mesoderm in our aggregoids, particularly in the double-mix condition, exhibited differentiation toward a sclerotome-like identity, a ventral somitic fate typically underrepresented in conventional gastruloids, potentially due to the absence of a localized Hedgehog (HH) signaling required for the somite ventral patterning (Hamazaki et al. 2024; Makwana et al. 2025; Tani et al. 2023). The prominent *IHH* and *SHH*-expressing midgut population in double-mix aggregoids could be a candidate for such a ventralizing signal source (Smirnova et al. 2025).

Together, our double and triple-mix aggregoid systems establish a reproducible modular platform for generating hSCBEMs. By combining distinct biased modules, this approach offers the potential to customize cell composition of the model to a specific research question and makes these models promising for investigating coordinated tissue co-development, with potential applications in developmental toxicology and drug screening. Our findings provide a framework for engineering lineage-biased hSCBEMs without prior genetic manipulation, expanding the experimental toolkit for studying early human development within an ethically informed research framework.

## Author contributions

I.M. and A.I. contributed equally. I.M., A.I. and S.K. conceived and designed the experiments; I.M., A.I., Y.B. and B.K.C. carried out the experiments; I.M., A.I., M.L., R.Z., S.P. and N.S. analyzed and elaborated data, M.L., T.M.L.A. and J.Ø. executed the scRNA-seq analysis, N.S., S.P. and E.M., N.G. provided critical comments to the manuscript; I.M., A.I., N.S. and S.K. wrote the manuscript. N.G. and S.K. provided funds.

## Acknowledgements

The project has received funding from the University of Oslo (UiO), the University Hospital of Oslo, the Research Council of Norway through its Centers of Excellence scheme grant Grant ID 262613, EU’s H2020 Marie Skłodowska-Curie Actions / COFUND Grant ID 801133, the European Innovation Council (EIC) (Horizon Europe) Pathfinder Challenge “Supervised Morphogenesis in Gastruloids” Grant ID 101071203 and the UIO:Life Science Convergence environments project ITOM.

## Declarations of interests

Stefan Krauss, Igor Meszka, Almira Ibragimova, Natalia Smirnova and Sergey Ponomartsev are named inventors on patent application NO20261501 with a priority date of 6 November 2026. The authors declare this intellectual property interest as a competing interest.

## Supplementary materials

**Supplementary Figure 1.**
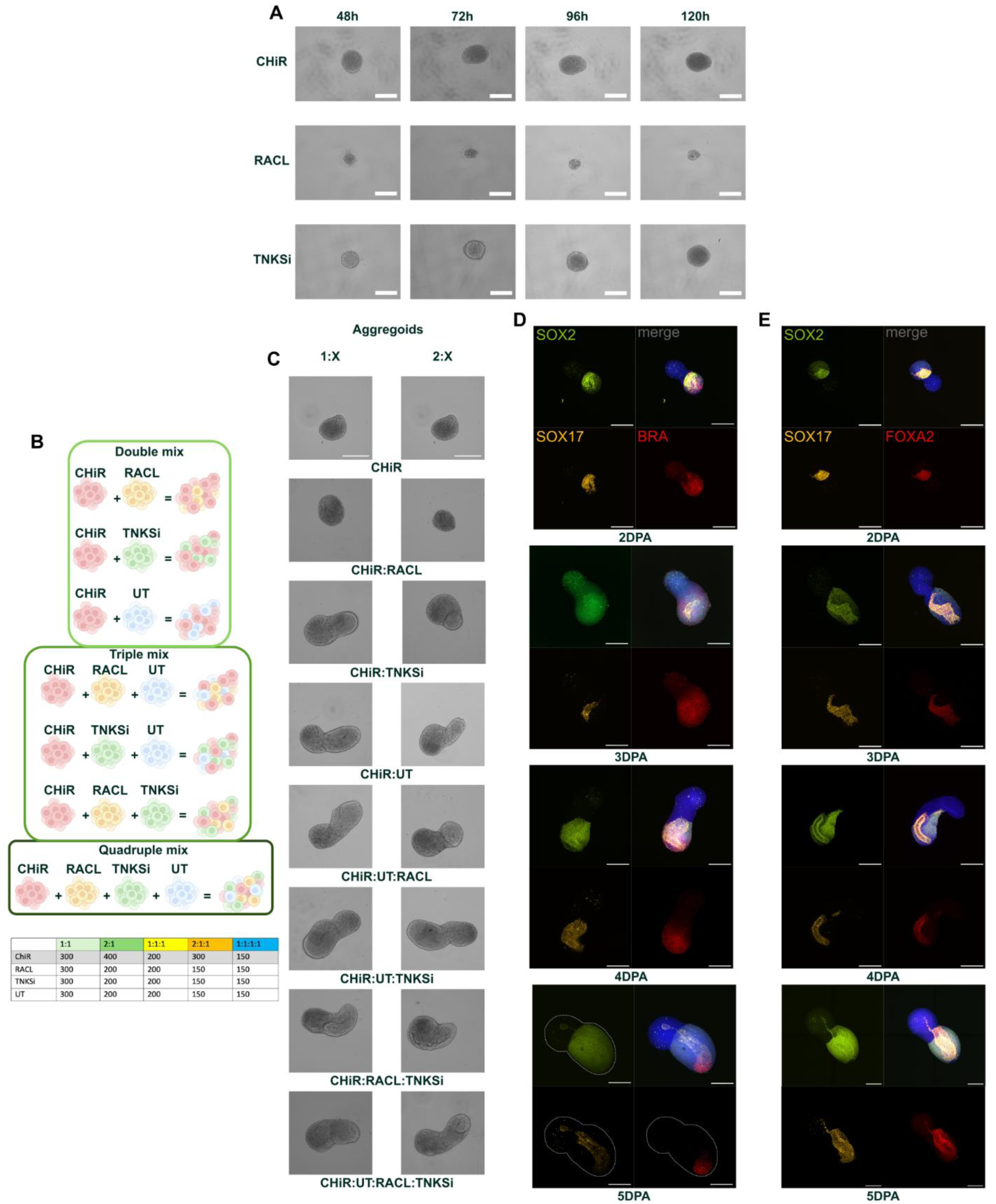
Pre-treatment creates fate-biased cell populations. **A)** Representative bright-field images of gastruloids created from the cells treated separately with CHIR, RACL and TNKSi across days post aggregation. Scale bar: 200 μm. **B)** Schematic representation of the tested mixes of the cells from different pre-treatments during the aggregation. **C)** Representative images of gastruloids phenotype depending on the mix and proportion of cells treated with ChiR, RACL, and TNKSi, as well as untreated cells. Images were taken on 5dpa. N=3 per condition. Scale bar: 300 μm. **D)** Immunofluorescence analysis of germ layer organization in day 5 gastruloids. SOX17 mark endodermal structures, SOX2 marks ectoderm, and BRA marks mesoderm, confirming spatial segregation and multi-lineage organization. Scale bar: 200 μm. **E)** Conformational immunofluorescence analysis of FOXA2 endodermal marker. Scale bar: 200 μm.

**Supplementary Figure 2.**
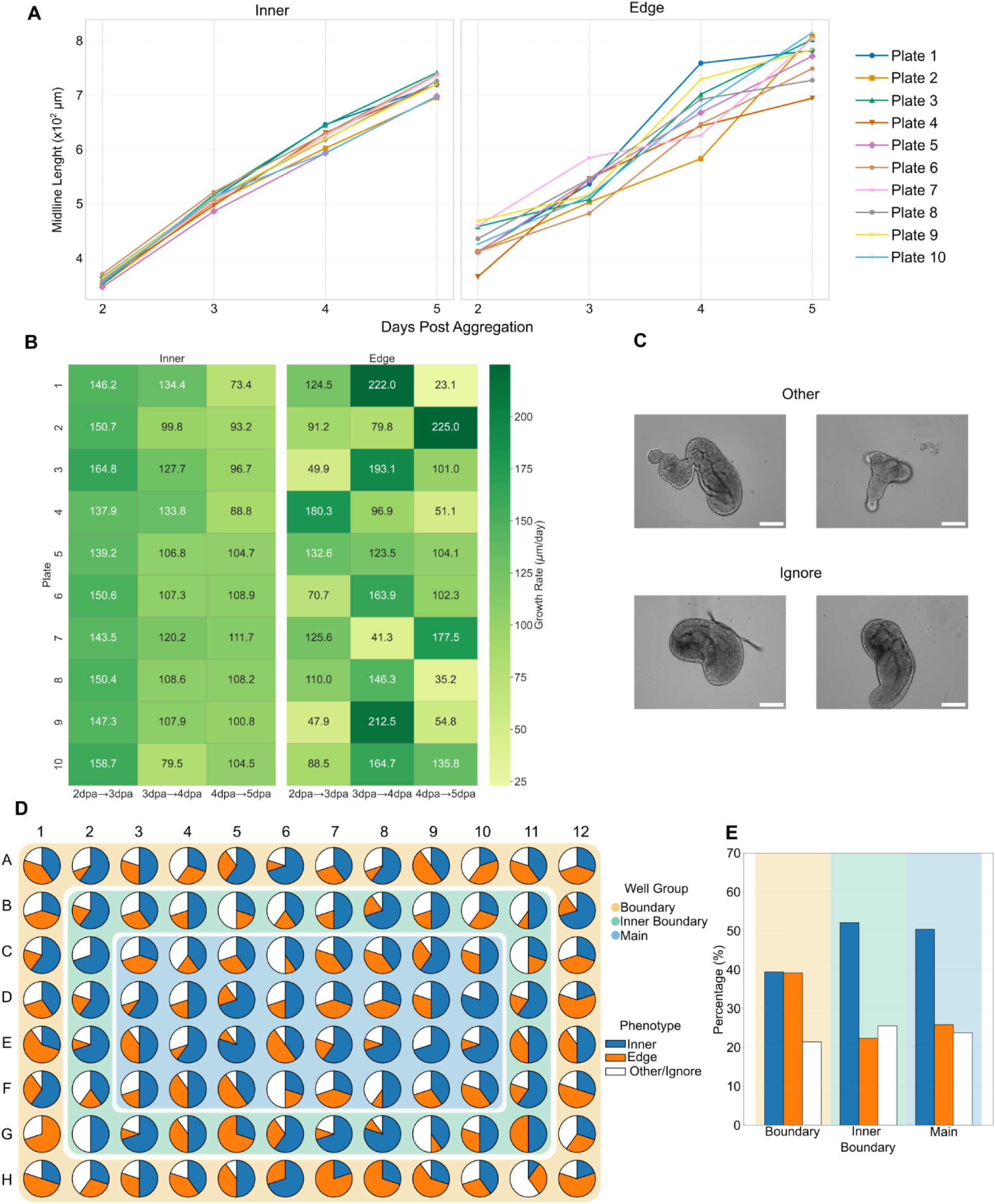
Co-aggregation strategies lead to emergence of reproducible hSCBEM morphotypes. **A)** Graph showing changes in midline length (eccentricity) depending on the day post aggregation and phenotype (I ner or Edge). n=10. **B)** Heatmap showing changes in growth rate across 10 plates depending on the day post aggregation and phenotype. n=10. **C)** Representative Brightfield images of the gastruloids which were not taken into the analysis. Scale bar: 200 μm. **D)** Schematic representation of the well group on the 96-well U-shaped plate. **E)** Graph showing the distribution of the different phenotypes depending on their location on the plate. n=3.

**Supplementary Figure 3.**
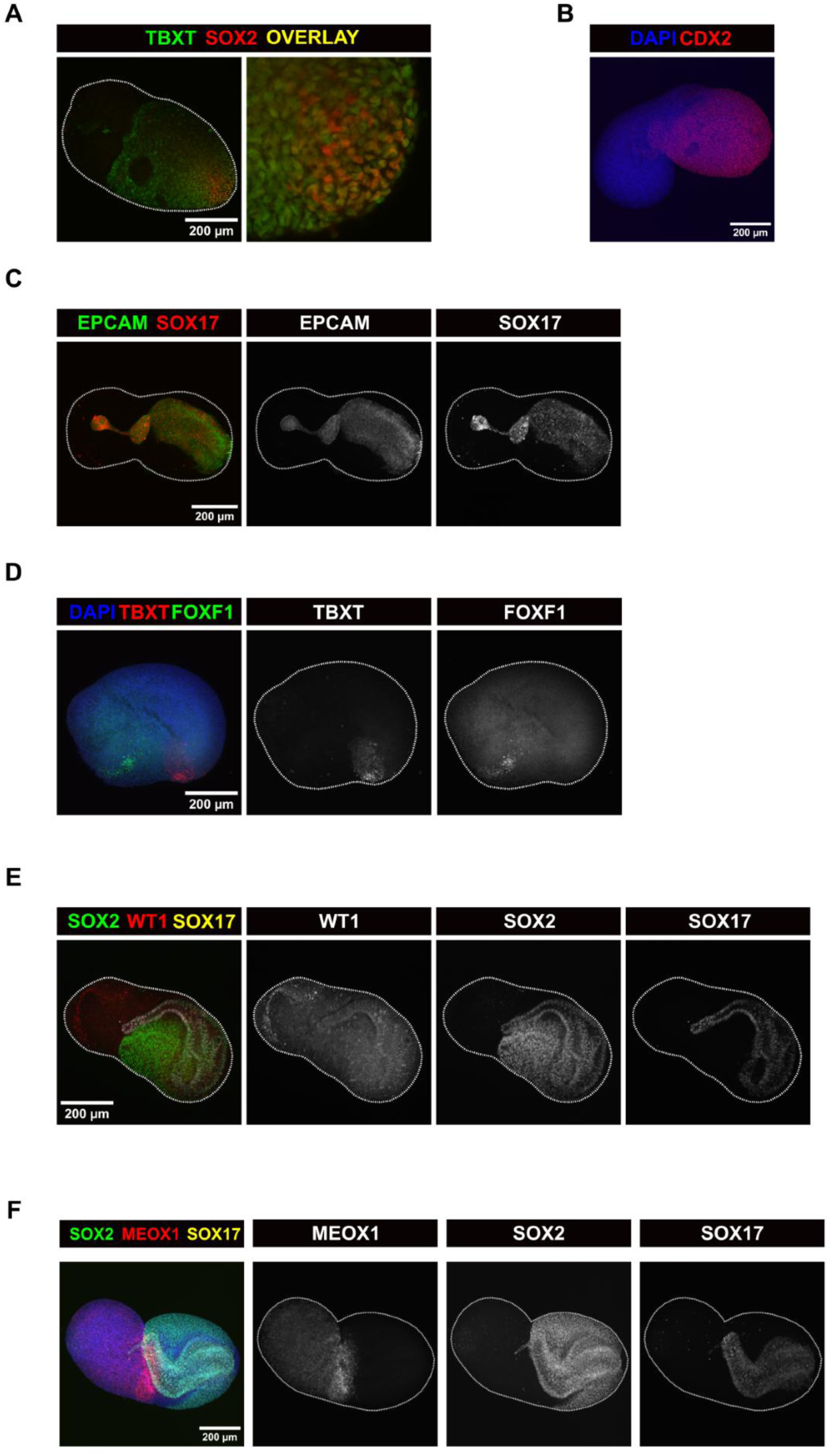
Single-cell transcriptomics reveal specific cell type diversification in hSCBEM. **A)** Representative confocal immunofluorescent image of TBXT, SOX2 colocalization in aggregoids at 120h. Scale bar: 200 μm **B)** Representative confocal immunofluorescent image of CDX2 localization in aggregoids at 120h. Scale bar: 200 μm **C)** Representative confocal immunofluorescent image of EPCAM and *SOX17*::mCherry colocalization in aggregoids at 120h. Scale bar: 200 μm **D)** Representative confocal immunofluorescent image of TBXT and FOXF1 expression in aggregoids at 120h. Scale bar: 200 μm **E)** Representative confocal immunofluorescent image of SOX2, WT1 and *SOX17*::mCherry expression in aggregoids at 120h. Scale bar: 200 μm **F)** Representative confocal immunofluorescent image of *SOX2*::mCitrine, MEOX1 and *SOX17*::mCherry expression in aggregoids at 120h. Scale bar: 200 μm

**Supplementary Figure 4.**
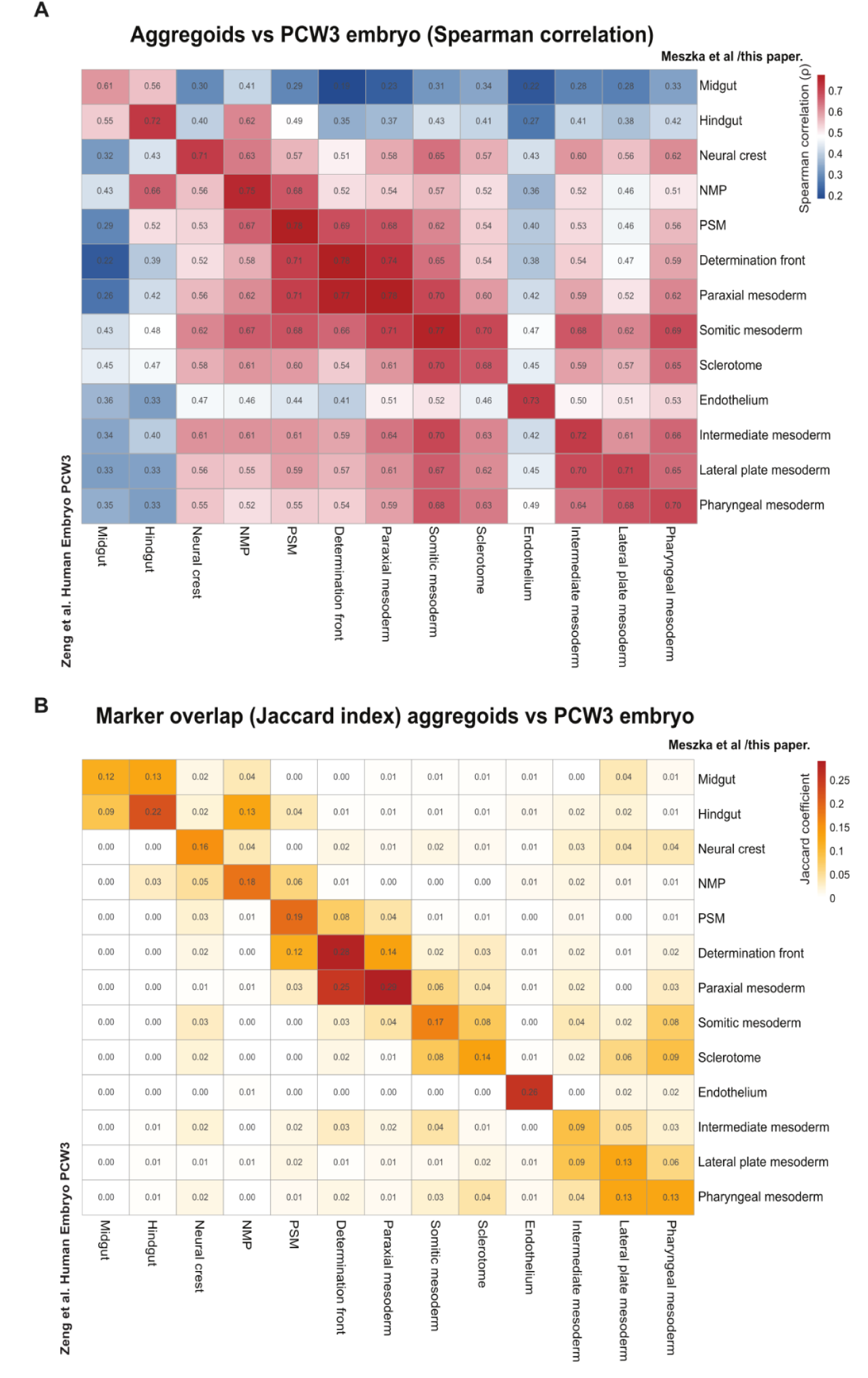
Transcriptional correspondence between aggregoid and human embryo cell types. Aggregoid cell types (columns) were compared with matched tissues of the PCW3 human embryo reference (Zeng et al., 2023) (rows), restricted to tissue types shared between the two datasets, using two complementary measures of transcriptional similarity. **(A)** Spearman correlation (ρ) between tissue-averaged transcriptomes, computed on the top 150 highly variable genes of the reference. **(B)** Marker overlap between matched tissues, quantified as the Jaccard index (shared markers / union) of the top 100 markers per tissue, ranked by average log2 fold-change; values range from 0 (no shared markers) to 1 (identical marker sets). Correspondence was robust to the gene and marker thresholds used, with comparable results across 100–500 genes.

## Methods

### Ethics

All experiments involving hSCBEM were conducted in accordance with 2021 ISSCR Guidelines for Stem Cell Research and Clinical Translation and the 2025 update on stem cell-based embryo models (SCBEMs). The project was approved by REK (Regional committees for medical and health research ethics). Case number 522684 from 01.06.2023. Cell lines used in this study were EU-registered and consented for use in such type of research.

### Human embryonic pluripotent stem cells culture

The following human embryonic stem cell (hESC) lines were used in this study: RUES2-GLR, obtained from Rockerfeller University (Martyn et al., 2018). hESCs were cultured in Essential 8 Medium (E8) (A1517001, Life Technologies) on 6-well plates coated with 1% (v/v) Geltrex (A1413302, Life Technologies) and maintained at 37°C in humidified atmosphere containing 5% CO₂. Culture medium was changed daily. Cells were passaged using 5 μM EDTA (T3924, Merck) in DPBS (14190-169, Life Technologies) upon reaching 60% confluency. All cell cultures were routinely screened for mycoplasma contamination every month using the MycoAlert Mycoplasma Detection Kit (LT07-318, Lonza). The karyotype of all cell lines was assessed annually to confirm genomic stability.

### Pre-treatment of hESC

Prior to co-aggregation, hESCs pools were exposed to one of three pre-treatment conditions. Cells were passaged at least 48h before treatment initiation. Untreated (UT) cells were kept in E8 (A1517001, Life Technologies) medium until co-aggregation. For the pre-treatment condition named CHIR, hESCs at approximately 60% confluency were exposed to 50 ng/mL Nocodazole (31430-18-9, Sigma-Aldrich) in E8 medium (A1517001, Life Technologies) for 6 h, beginning 30 h prior to co-aggregation. Following Nocodazole treatment, cells were washed three times with DPBS (14190-169, Life Technologies). At 24 h prior to co-aggregation, the medium was replaced with E8 medium supplemented with 6 μM CHIR99021 (4423, Tocris Bioscience), and cells were maintained under these conditions until aggregation. In the RACL pre-treatment condition, hESCs at approximately 30% confluency were cultured in RACL medium for 48 h prior to co-aggregation. RACL medium consisted of RPMI 1640 (11875093, Gibco) supplemented with 1× B27 Supplement Minus Insulin (A1895601, Gibco), 100 ng/mL Activin A (120-14P, PeproTech), 3 μM CHIR99021 (4423, Tocris Bioscience), and 10 ng/mL human LIF (300-05, PeproTech). Medium was refreshed every 24 h. For TNKSi condition, Tankyrase 1/2 inhibitor OM-153 (Waaler et al., 2020) in E8 (A1517001, Life Technologies) was administered at a concentration of 1 µM for 48 hours prior to aggregation to the hESCs at approximately 30% confluency.

### Generation of aggregoids

Prior to aggregation, pre-treated hESCs were washed 2-3 times with DPBS (14190-169, Life Technologies) and dissociated into a single-cell suspension using Accutase (A6964, Merck). Cells were centrifuged at 100 g for 4 min and resuspended in E6 medium (A1516401, Life Technologies) supplemented with 1 μM CHIR99021 (4423, Tocris Bioscience), 2 μM SB431542 (616461, Sigma-Aldrich), and 5 μM Y-27632 (72304, STEMCELL Technologies). Cell numbers were counted using a hemocytometer. Pre-treated hESCs were resuspended in 5 mL of aggregation medium and seeded into ultra-low attachment 96-well plates (650970, Greiner Bio-One) at a density of 600 cells (either 200 cells of each pretreatment in triple or 300 cells of each in double) per well in a final volume of 40 μL per well. Following seeding, plates were centrifuged at 200 g for 2 min to facilitate cell aggregation and subsequently incubated at 37°C in a humidified atmosphere containing 5% CO₂. Aggregates were cultured under these conditions for 24 h. 24h after aggregation, 160 μL of E6 medium (A1516401, Life Technologies) was added to each well. From day 48h post-aggregation onwards, 100 μL of medium was replaced daily with fresh E6 (A1516401, Life Technologies) medium.

### Immunostaining

The hSCBEMs that were collected for immunostaining were first washed in PBS (supplemented with MgCl₂ and CaCl₂) containing 0.5% BSA. They were then fixed with 4% PFA for 30 minutes at room temperature. The hSCBEMs were then washed three times with PBS. Next, hSCBEMs were permeabilised with PBST (PBS supplemented with 0.05% Triton X-100) for 20 minutes at room temperature, then blocked overnight in blocking solution (PBS + 0.05% Triton X-100 + 10% FBS). hSCBEMs were then incubated with the primary antibody in the blocking solution at 4 °C for 72–96 hours. After this time, hSCBEMs were washed three times with PBST and left in the blocking solution overnight. Secondary antibodies in the blocking solution were then added to the gastruloids, which were incubated for 24 hours at 4 °C. After incubation with the secondary antibodies, hSCBEMs were washed three times with the blocking solution and once with PBST. The sample was fixed once again in 4% PFA for 20 minutes. hSCBEMs were then washed twice in 0.02 M phosphate buffer (25 mM NaH₂PO₄ + 75 mM Na₂HPO₄). The samples were then cleared with a clearing agent overnight.

### Preparation and fixation of single-cell suspension for single-cell RNA sequencing

Cells from four 2D culture conditions (UT, CHIR, RACL, and TNKSi) and three hSCBEM conditions (CHIR:RACL, CHIR:RACL:TNKSi (Edge), and CHIR:RACL:TNKSi (Inner)) were collected at 48 h, 72 h, 96 h, and 120 h. For hSCBEM samples, 384 aggregates were pooled at 48 h, 288 aggregates at 72 h, 192 aggregates at 96 h, and 96 aggregates at 120 h. To prepare single-cell suspension, the collected hSCBEM were transferred to a 200 μl drop of TrypLE (12604013, Gibco) and incubated at 37 °C with pipetting every 5 minutes until complete dissociation. Subsequently, the TrypLE was quenched with 800 μl of ice-cold DPBS (14190-169, Life Technologies) containing 0.5% BSA (A8412, Merck). The cell suspension was centrifuged at 4 °C and 300 g for 5 minutes after filtering through a cell strainer (STEMCELL Technologies). After removal of the supernatant, the cell pellet was washed twice with 1 mL of DPBS supplemented with 0.04% BSA, centrifuging at 300 g for 5 min at 4°C. Following the final wash, the cell pellet was resuspended in 1 mL of DPBS containing 0.04% BSA. The supernatant was then discarded and the cell pellet washed with 1 ml of PBS + 0.04% BSA, followed by centrifugation at 4°C and 300 g for 5 minutes. This process was repeated once more, after which the cell pellet was dissociated in 1 ml of DPBS + 0.04% BSA (A8412, Merck). Cell viability was assessed by Trypan Blue (15250-061, Gibco) staining. Amount of dead cells was less than 5% for all samples. Cells fixation was done using Chromium Next GEM Single Cell Fixed RNA Sample Preparation Kit (16 reactions; 10X Genomics, Cat. No. 1000414) and stored at -80°C or 4°C until library preparation.

### Library Preparation and Single-Cell RNA Sequencing

Samples were multiplexed, and cDNA libraries were generated from 10,000 cells per sample using the Chromium Fixed RNA Profiling Kit (1000496, 10x Genomics) according to the manufacturer’s instructions. Library quality and concentration were assessed using the High Sensitivity DNA Kit on an Agilent 2100 Bioanalyzer (Agilent Technologies). Sequencing was performed at the Norwegian Sequencing Centre on a NovaSeq X Plus platform (NovaSeq X Plus System Suite v1.2.2.48004; Illumina). Libraries were sequenced on a single 25B flow-cell lane using a 300-cycle sequencing kit.

### Data normalization

The sequenced data was aligned against the human GRCh38 reference genome using the 10x Genomics Cell Ranger (v8.0.1) pipeline and the Chromium Human Transcriptome Probe Set (v1.0.1), and de-multiplexed in silico accordingly. The samples were then aggregated in the same pipeline, with normalization and batch-correction, or used non-aggregated where applicable.

### Preprocessing and Quality control

The count matrices generated by the CellRanger pipeline (v8.0.1) were loaded into R (v4.5.1) using the Seurat library (v5.3.0). Raw count matrices were imported per sample with Read10X_h5 and converted to Seurat objects. Cells were retained if they had <200 and ≤7,500 detected genes (nFeature_RNA), <400 and ≤40,000 UMIs (nCount_RNA). The mitochondrial content of each cell was computed from MT-genes (PercentageFeatureSet), and cells with ≥40% mitochondrial reads were removed. Each sample was log-normalized (NormalizeData, scale factor 10,000), and the 2,000 most variable genes were selected (FindVariableFeatures, vst). Data were scaled, and principal component analysis (PCA) was run on the variable features. Cells were clustered using the first 15 principal components (FindNeighbors, FindClusters, resolution 0.5) and embedded with UMAP (15 PCs). Cell-cycle phase scores were assigned with CellCycleScoring (Seurat S- and G2M-phase gene sets) and regressed out during scaling (vars.to.regress = S.Score, G2M.Score). Per-sample diagnostic plots were inspected, and clusters identified as low quality were removed manually on a per-sample basis. The full normalization–clustering pipeline was then re-run on the filtered objects.

### Doublet detection and removal

Doublets were identified per sample with scDblFinder, run on the SingleCellExperiment representation of each object. Cells with a doublet score ≥ 0.7 were classified as high-confidence doublets and removed; lower-confidence calls were retained. The standard normalization, cell-cycle regression, clustering and UMAP pipeline was re-run on the singlet-filtered objects, yielding final high-quality cells.

### Data Integration

The filtered, singlet objects were integrated using reciprocal PCA (rPCA) in Seurat. Two complementary integration strategies were applied:

- By condition (condition_2_1 - double, condition_3_1 - inner, condition_3_2 - edge): the four time points sharing a treatment condition were integrated
- By day (day2–day5): the three conditions collected on the same day were integrated

For each integration set, objects were normalized and variable features identified, then 5,000 integration features were selected (SelectIntegrationFeatures). Anchors were found with FindIntegrationAnchors (reduction = “rpca”, 50 dimensions) and the data integrated with IntegrateData (50 dimensions). The integrated assay was scaled, followed by PCA (50 components), graph-based clustering (50 PCs, resolution 0.5) and UMAP (50 PCs). Cluster marker genes were identified with FindAllMarkers on the RNA assay (only.pos = TRUE, min.pct = 0.25, logfc.threshold = 0.25).

### Cell-type annotation

#### Initial annotation

Day-integrated aggregoid objects were first annotated using curated marker gene sets for endothelial, gut endoderm, intermediate mesoderm, lateral-plate-mesoderm-like/early pharyngeal, paraxial mesoderm, presomitic mesoderm (PSM) and somite identities. For each gene set, per-cell module scores were computed with UCell (AddModuleScore_UCell). Each cell was assigned the identity of its highest-scoring module; cells whose maximum module score did not exceed 0.1 were labelled “Unknown”. Marker expression was visualised with feature plots, dot plots, violin plots and ridge plots. These assignments provided a scaffold that was refined as described below.

#### Reference embryo

Cells of the PCW3 embryo were annotated by refining the published Zeng et al. (2023) labels through systematic, marker-based curation. Candidate identities were evaluated on canonical lineage markers using feature-resolved UMAPs and by module scoring (AddModuleScore) of lineage-specific gene sets; cells misassigned within clusters were manually reassigned by interactive selection on the embedding (CellSelector), and ambiguous populations were cross-checked against a mouse embryo reference atlas (Imaz-Rosshandler et al., 2024). This procedure resolved the published annotation (19 categories) into 36 biologically specific cell types spanning all three germ layers, subdividing broader mesodermal, neural, endodermal and haematopoietic groups into their constituent lineages.

#### Aggregoids and gastruloids

Both the aggregoid and gastruloid datasets were subsequently annotated independently using the same marker- and module-score-driven strategy. Candidate lineage programmes were scored per cluster (AddModuleScore), and identities were assigned only when supported concordantly by co-expression frequency (the fraction of double- or triple-positive cells), per-cluster marker ranking and marker-specificity checks. Finer populations were resolved with FindSubCluster and, where required, by manual curation on the UMAP. Gut endoderm was regionalized into foregut, midgut, hindgut and early gut-endoderm identities by tiered module scoring against anteroposterior gut markers. Module scores driven by non-specific genes were identified and excluded (for example, a cardiomyocyte score inflated by titin (*TTN*) expression in skeletal muscle was corrected to a heart-restricted marker set).

#### Cross-dataset label harmonization

For joint analysis, the annotations of the three datasets were reconciled into a shared label vocabulary. In all reported embeddings, each dataset is coloured by its own harmonized annotation.

### Single-cell RNA-seq datasets and quality control

Three single-cell RNA-sequencing datasets were integrated: a post-conceptional week 3 (PCW3; Carnegie stage 10) human embryo as the reference (Zeng et al., 2023; GEO: GSE155121), and two human gastruloid datasets as queries — the human gastruloids of Hamazaki et al. (2024; GEO: GSE208369) and our in-house aggregoid model (Days 2– 5). All processing was performed in R using Seurat v5 (Hao et al., 2024). For each dataset, a count matrix was used to construct a Seurat object and curated per-cell annotations were joined to cells by barcode. Following doublet removal, reference cells were retained if they expressed more than 2,500 genes and fewer than 10% mitochondrial reads; both gastruloid datasets were filtered further, retaining cells with more than 1,500 detected genes. This threshold was applied in addition to the per-sample quality control described above, and only for the cross-dataset comparison. The Hamazaki dataset was restricted to timepoints from 48 h onward (48, 72 and 96 h conventional; 96 and 120 h retinoic acid) for comparison to the same time points in the aggregoid dataset. Counts were log-normalized, the 3,000 most variable genes were identified by the variance-stabilizing transformation, and the data were scaled prior to principal-component analysis (70 components).

### Reference mapping

Aggregoid cells at 96 h (n = 9,476) and 120 h (n = 10,073), conventional gastruloids at 96 h (n = 8,018) and RA-treated gastruloids at 96 h and 120 h (n = 1,195 and 9,988) were mapped separately onto the reference. A UMAP model was computed for the reference on principal components 1–68 (n.neighbors = 30, min.dist = 0.3, seed = 42). For each query, transfer anchors were identified by projecting log-normalized query data onto the reference PCA (FindTransferAnchors, dimensions 1–68). Cell-type labels were transferred and query cells projected onto the reference UMAP using MapQuery (k.weight = 50) (Stuart et al., 2019; Hao et al., 2021, 2024). Aggregoid conditions (Double, Triple Inner, Triple Edge) were mapped together within each timepoint and separated afterwards for display.

### Marker-based checks of transferred labels

Labels for three groups of cell types that are easily confused with neighbouring types were also checked against marker genes: brain regions (forebrain, midbrain, hindbrain) against neural-biased NMPs, neural crest and spinal cord; placodal ectoderm against gut endoderm (foregut, midgut, hindgut); and pharyngeal mesoderm and cranial mesenchyme against anterior and trunk lateral plate mesoderm. For each of these, we used the reference to find the genes that best distinguish it from other tissues (the top 25 differentially expressed genes in each direction). Cells were then scored on how strongly they expressed the cell type’s genes relative to the other cell types. A transferred label was kept only if the cell scored at least as high as the lowest-scoring 10% of genuine reference cells of that type.

Because the reference contained few gut-endoderm cells, cells expressing the pan-endoderm genes *SOX17, FOXA2* and *FOXA1* at reference gut levels were assigned to gut endoderm. Low-confidence predictions were kept when the uncertainty was only between closely related subtypes of the same lineage (for example, somitic and paraxial mesoderm). Pharyngeal mesoderm labels were kept only when supported by pharyngeal markers other than TBX1, which may also be expressed in lateral plate mesoderm as a response to hedgehog signalling.

All criteria were applied identically to the aggregoid and gastruloid datasets.

### Visualization and composition analysis

In the reference projections (Figure 4), query cells are shown in the reference UMAP embedding and coloured by their gated label; The reference contains too few gut cells for regional label transfer, so gut regional identity (midgut, hindgut, gut endoderm) in query panels is taken from each dataset’s own annotation. Cell-type proportions across systems (Figure 4) were calculated from each dataset’s harmonized annotation as the percentage of cells per system, excluding cells flagged as artefacts.

### Transcriptional correspondence between aggregoid and embryo cell types

For cell types shared between the two datasets, tissue-averaged expression profiles were computed across all cells of each annotated type. Transcriptional similarity was quantified by Spearman correlation (ρ) across the top 150 highly variable genes of the PCW3 reference and visualised as a heatmap (Supplementary Figure 4). Correspondence was robust to this threshold, with comparable results obtained across 100–500 genes. In parallel, marker overlap was quantified as the Jaccard index (shared markers / union of markers) of the top 100 markers per tissue, ranked by average log2 fold-change and identified with FindAllMarkers. Both measures were visualised as heatmaps (Supplementary Figure 4A, B). Correspondence was robust to these thresholds, with comparable results obtained across 100–500 genes.

### Software

Analyses were performed in R v4.5.1 with Seurat v5.3.0, SeuratObject, UCell, scDblFinder, DESeq2, apeglm, ggplot2 v4.0.3, patchwork, ggrepel, Polychrome and ggrastr.

### Data management

All scRNAseq data generated during this study were stored and managed using the TSD (Tjenester for Sensitive Data) facilities, owned by the University of Oslo and developed by the TSD service group at the University of Oslo IT Department. TSD provides a secure computing environment for the storage, processing, and analysis of sensitive research data in accordance with institutional and national data protection requirements.

### Imaging

Automated brightfield and fluorescence imaging of hSCBEMs was performed using an EVOS M7000 imaging system (Thermo Fisher Scientific). Brightfield images were acquired at the indicated time points between 48 h and 120 h post-aggregation using a 4x or 10× objective. Fluorescence imaging of *SOX17*::mCherry and *SOX2::*mCitrine reporter lines was performed using RFP (AMEP4952, Thermo Fisher Scientific) and YFP (AMEP4954, Thermo Fisher Scientific) light cubes, respectively. Fixed and stained structures were imaged using a Crest Spinning disk (X-light V3 CREST). Z-stack images were acquired with a z-step of 3 μm. For confocal imaging, hSCBEMs were mounted in RAPICLEAR (152001, SunJin Lab).

### Midline related measurements

The midline extraction workflow was adapted from the algorithm published by Merle et al. in 2024. Each hSCBEMs image set consisted of three channels (*SOX17*::mCherry, *SOX2*::mCitrine and grayscale), which were pre-processed into a stacked RGB ‘.tiff’ file to optimize computational loading while maintaining the original pixel resolution and intensity. Channels were arranged as: red (R) – *SOX17*::mCherry, green (G) – *SOX2*::mCitrine, blue (B) – grayscale. The midline profile measurement procedure consisted of four major steps: сontour extraction where the hSCBEMs boundary was identified from the grayscale image channel, and this contour mask was subsequently applied to the *SOX17*::mCherry and *SOX2*::mCitrinechannels to ensure that all downstream measurements were performed within the biologically meaningful region. Skeleton extraction: a centreline was obtained by skeletonizing the extracted gastruloid boundary. The skeleton was then ordered and extended to reach both terminal ends of hSCBEMs. Polarization: hSCBEMs polarity was determined based on the Sox2 intensity distribution along the midline. The midline end region exhibiting higher Sox2 intensity was considered posterior with the opposite end defined as anterior. Slicing and normalization: to quantify fluorescence intensity variation along the anterior-posterior axis, each hSCBEMs was sliced along midline into 21 equally spaced segments. For each segment, channel-specific intensity profiles, pixel area and corresponding signal statistics were extracted for quantitative analysis. To enable meaningful comparison across hSCBEMs of different sizes, intensity profiles were normalized by the area of each segment, ensuring that fluorescence measurements were not biased by geometric differences or segmentation variability.

### CellProfiler Measurements

CellProfiler is a widely used software for pixel based image analysis and quantitative morphological analysis (Stirling et al., 2021). Version 4.2.4 and 4.2.8 were used in this study, with identical analysis pipelines applied across all datasets. To characterize gastruloid morphology and growth dynamics across time points, CellProfiler measurements were applied to segment and quantify all images from day 2 to day 5, each containing the three channels (*SOX17*::mCherry, *SOX2*::mCitrine, grayscale). From each channel, three biologically relevant object types were extracted: red primary object: high intensity regions in the *SOX17*::mCherry channel; green primary object: high intensity regions in the Sox2 channel; grayscale object: regions occupied by the actual hSCBEMs in the grayscale channel. For each object type, a comprehensive set of morphological and pixel-based features was quantified, including: area metrics (object size, bounding box), shape metrics (eccentricity, solidity), intensity features (mean, median intensity), texture features (spatial variation in pixel intensity), colocalization (quantify overlap between different channels).

## References

Anlas, K., Nakaki, F., Gritti, N., Font-Reverter, J., Arató, K., Planasdemunt-Hospital, M., Oriola, D., Sharpe, J., & Trivedi, V. (2025). Axial patterning of gastruloids via diverse cell type compositions. bioRxiv. 10.1101/2025.03.26.645494

Andreatta, M., and Carmona, S.J. (2021). UCell: robust and scalable single-cell gene signature scoring. Comput. Struct. Biotechnol. J. 19, 3796–3798.

Ayyappan, V., Triandafillou, C. G., Sarma, K., & Raj, A. (2026). Gastruloid patterning reflects division of labor among biased stem cell clones. bioRxiv. 10.1101/2025.07.12.664536

Azami, T., Patton, E. E., & Nichols, J. (2026). Simplified in vitro generation of human gastruloids for modelling early development. Development, 153(20), dev205719. 10.1242/dev.205719

Beccari, L., Moris, N., Girgin, M., Turner, D. A., Baillie-Johnson, P., Cossy, A. C., Lutolf, M. P., Duboule, D., & Martinez-Arias, A. (2018). Multi-axial self-organization properties of mouse embryonic stem cells into gastruloids. Nature, 562(7726), 272–276. 10.1038/s41586-018-0578-0

Bentzen, H. B., Gaillard, M., Nachman, I., Reumann, D., Gadegaard, N., David, L., Lanner, F., Moris, N., Pasque, V., Rivron, N., Sozen, B., Isasi, R., Krauss, S., & Veenvliet, J. V. (2026). A guide to using embedded ethics in human stem-cell-based embryo model research. Nature Cell Biology, 28, 665–673. 10.1038/s41556-026-01909-9

Blotenburg, M., Suurenbroek, L., Bax, D., de Visser, J., Bhardwaj, V., Braccioli, L., de Wit, E., van Boxtel, A., Marks, H., & Zeller, P. (2025). Stem cell culture conditions affect in vitro differentiation potential and mouse gastruloid formation. PLoS ONE, 20, e0317309.

Cermola, F., D’Aniello, C., Tate, R., De Cesare, D., Martinez-Arias, A., Minchiotti, G., & Patriarca, E. J. (2021). Gastruloid development competence discriminates different states of pluripotency. Stem Cell Reports, 16, 354–369. 10.1016/j.stemcr.2020.12.013

Chen, G., Gulbranson, D. R., Hou, Z., Bolin, J. M., Ruotti, V., Probasco, M. D., Smuga-Otto, K., Howden, S. E., Diol, N. R., Propson, N. E., Wagner, R., Lee, G. O., Antosiewicz-Bourget, J., Teng, J. M. C., & Thomson, J. A. (2011). Chemically defined conditions for human iPSC derivation and culture. Nature Methods, 8(5), 424–429. 10.1038/nmeth.1593

Cliff, T. S., Wu, T., Boward, B. R., Yin, A., Yin, H., Glushka, J. N., Prestegaard, J. H., & Dalton, S. (2017). MYC controls human pluripotent stem cell fate decisions through regulation of metabolic flux. Cell Stem Cell, 21(4), 502–516.e9. 10.1016/j.stem.2017.08.018

Das, A., Patil, S., Ravi, K., & Inamdar, M. S. (2026). A robust, reproducible, accessible and scalable protocol for generating three-dimensional human gastruloids. bioRxiv. 10.1101/2026.05.23.727089

Dias, A., Pascual-Mas, P., Robertson, G., Torregrosa-Cortés, G., Stelloo, S., Casaní-Galdón, P., Babin, S., Romaniuk, Y., Mayran, A., Wehmeyer, A. E., Garcia-Ojalvo, J., McNamara, H. M., Vermeulen, M., Arnold, S. J., & Martinez-Arias, A. (2025). Opposing Nodal and Wnt signalling activities govern the emergence of the mammalian body plan. bioRxiv, 2025.01.11.632562. 10.1101/2025.01.11.632562

Dingare, C., & Steventon, B. (2023). Gastruloids—A minimalistic model to study complex developmental metabolism. Emerging Topics in Life Sciences, 7(4), 455–464. 10.1042/ETLS20230082

Dingare, C., Cao, D., Yang, J. J., Sozen, B., & Steventon, B. (2024). Mannose controls mesoderm specification and symmetry breaking in mouse gastruloids. Developmental Cell, 59(12), 1523–1537.e6. 10.1016/j.devcel.2024.03.031

Farag, N., Sacharen, C., Avni, L., & Nachman, I. (2024). Coordination between endoderm progression and mouse gastruloid elongation controls endodermal morphotype choice. Developmental Cell, 59, 2364– 2374.e2364.

Germain, P.-L., Lun, A., Garcia Meixide, C., Macnair, W., and Robinson, M.D. (2022). Doublet identification in single-cell sequencing data using scDblFinder. F1000Res. *10*, 979. 10.12688/f1000research.73600.2

Hamazaki, N., Yang, W., Kubo, C. A., Qiu, C., Martin, B. K., Garge, R. K., Regalado, S. G., Nichols, E. K., Pendyala, S., Bradley, N., Fowler, D. M., Lee, C., Daza, R. M., Srivatsan, S., & Shendure, J. (2024). Retinoic acid induces human gastruloids with posterior embryo-like structures. Nature Cell Biology, 26(10), 1790– 1803. 10.1038/s41556-024-01487-8

Hashmi, A., Tlili, S., Perrin, P., Lowndes, M., Peradziryi, H., Brickman, J. M., Martínez-Arias, A., & Lenne, P.-F. (2022). Cell-state transitions and collective cell movement generate an endoderm-like region in gastruloids. eLife, 11, e59371. 10.7554/eLife.59371

Hao, Y., Hao, S., Andersen-Nissen, E., Mauck, W.M., 3rd, Zheng, S., Butler, A., Lee, M.J., Wilk, A.J., Darby, C., Zager, M., et al. (2021). Integrated analysis of multimodal single-cell data. Cell 184, 3573– 3587.e29. 10.1016/j.cell.2021.04.048

Hao, Y., Stuart, T., Kowalski, M.H., Choudhary, S., Hoffman, P., Hartman, A., Srivastava, A., Molla, G., Madad, S., Fernandez-Granda, C., et al. (2024). Dictionary learning for integrative, multimodal and scalable single-cell analysis. Nat. Biotechnol. 42, 293–304. 10.1038/s41587-023-01767-y

Imaz-Rosshandler, I., Rode, C., Guibentif, C., Harland, L.T.G., Ton, M.-L.N., Dhapola, P., et al. (2024). Tracking early mammalian organogenesis – prediction and validation of differentiation trajectories at whole organism scale. Development 151, dev201867. 10.1242/dev.201867

Laco, F., Woo, T. L., Zhong, Q., Szmyd, R., Ting, S., Khan, F. J., Chai, C. L., Reuveny, S., Chen, A., & Oh, S. (2018). Unraveling the inconsistencies of cardiac differentiation efficiency induced by the GSK3β inhibitor CHIR99021 in human pluripotent stem cells. Stem Cell Reports, 10(6), 1851–1866. 10.1016/j.stemcr.2018.03.023

Linneberg-Agerholm, M., Wong, Y. F., Herrera, J. A. R., Monteiro, R. S., Anderson, K. G. V., & Brickman, J. M. (2019). Naïve human pluripotent stem cells respond to Wnt, Nodal and LIF signalling to produce expandable naïve extra-embryonic endoderm. Development, 146(24). 10.1242/dev.180620

Liu, L., & Wu, J. (2025). Stem cell-based models of early human development. Development, 152.

Makwana, K., Tilley, L., Chakravarty, P., et al. (2025). Modelling co-development between the somites and neural tube in human trunk-like structures. Nature Cell Biology, 27(12), 2049–2062. 10.1038/s41556-025-01813-8

Martinez-Arias, A., Dias, A., Wehmeyer, A. E., Arnold, S. J., & Fiuza, U. M. (2025). A modular organization of mammalian gastrulation and the Spemann-Mangold organizer. Cells & Development, 204031.

Martinez-Arias, A., Marikawa, Y., & Moris, N. (2022). Gastruloids: Pluripotent stem cell models of mammalian gastrulation and embryo engineering. Developmental Biology, 488, 35–46.

Martinez-Arias, A., Rivron, N., Tajbakhsh, S., Johnston, J., Alev, C., Bally-Cuif, L., Böke, E., Cavazza, T., Cave, E., Cyranoski, D., David, L., Esteban, M. A., Fu, J., Geijsen, N., De Graeff, N., Hanna, J. H., Hopwood, N., Inamdar, M. S., Lanner, F., … Yu, L. (2026). Human stem cell-based embryo models: Innovation, ethics, and policy. Human Reproduction, 41(6), 830–846. 10.1093/humrep/deag035

Martyn, I., Kanno, T. Y., Ruzo, A., Siggia, E. D., & Brivanlou, A. H. (2018). Self-organization of a human organizer by combined Wnt and Nodal signalling. Nature, 558(7708), 132–135. 10.1038/s41586-018-0150-y

McNamara, H. M., Solley, S. C., Adamson, B., Chan, M. M., & Toettcher, J. E. (2024). Recording morphogen signals reveals mechanisms underlying gastruloid symmetry breaking. Nature Cell Biology, 26(11), 1832–1844. 10.1038/s41556-024-01521-9

Merle, M., Friedman, L., Chureau, C., Shoushtarizadeh, A., and Gregor, T. (2024). Precise and scalable self-organization in mammalian pseudo-embryos. Nat. Struct. Mol. Biol. 31, 896–902. 10.1038/s41594-024-01251-4

Miao, Y., Djeffal, Y., De Simone, A., Zhu, K., Lee, J. G., Lu, Z., Silberfeld, A., Rao, J., Tarazona, O. A., Mongera, A., Rigoni, P., Diaz-Cuadros, M., Song, L. M. S., Di Talia, S., & Pourquié, O. (2023). Reconstruction and deconstruction of human somitogenesis in vitro. Nature, 614, 500–508.

Moris, N., Anlas, K., van den Brink, S. C., Alemany, A., Schröder, J., Ghimire, S., Balayo, T., van Oudenaarden, A., & Martinez-Arias, A. (2020). An in vitro model of early anteroposterior organization during human development. Nature, 582(7812), 410–415. 10.1038/s41586-020-2383-9

Moris, N., Martinez-Arias, A., Pera, M., Rivron, N., Sermon, K., & de Graeff, N. (2026). Morally relevant features warranting ethical oversight in human stem cell-based embryo models. Nature Cell Biology, 28, 659–664.

Neupane, J., Lubatti, G., Gross-Thebing, T., Ruiz Tejada Segura, M. L., Butler, R., Gross-Thebing, S., Dietmann, S., & Surani, M. A. (2025). The emergence of human primordial germ cell-like cells in stem cell-derived gastruloids. Science Advances, 11, eado1350.

Olmsted, Z. T., & Paluh, J. L. (2021). Co-development of central and peripheral neurons with trunk mesendoderm in human elongating multi-lineage organized gastruloids. Nature Communications, 12(1), 3020. 10.1038/s41467-021-23294-7

Olmsted, Z. T., & Paluh, J. L. (2022). A combined human gastruloid model of cardiogenesis and neurogenesis. iScience, 25(6), 104486. 10.1016/j.isci.2022.104486

Olmsted, Z. T., Paredes-Espinosa, M. B., & Paluh, J. L. (2022). Generation of human elongating multi-lineage organized cardiac gastruloids. STAR Protocols, 3.

Pera, M. F., de Wert, G., Dondorp, W., Lovell-Badge, R., Mummery, C. L., Munsie, M., & Tam, P. P. (2015). What if stem cells turn into embryos in a dish? Nature Methods, 12, 917–919.

Rivron, N. C., Frias-Aldeguer, J., Vrij, E. J., Boisset, J., Korving, J., Vivié, J., Truckenmüller, R. K., Van Oudenaarden, A., Van Blitterswijk, C. A., & Geijsen, N. (2018). Blastocyst-like structures generated solely from stem cells. Nature, 557(7703), 106–111. 10.1038/s41586-018-0051-0

Rivron, N. C., Martinez-Arias, A., Pera, M. F., Moris, N., & M’hamdi, H. I. (2023). An ethical framework for human embryology with embryo models. Cell, 186(17), 3548–3557. 10.1016/j.cell.2023.07.028

Rossant, J., & Tam, P. P. (2022). Early human embryonic development: Blastocyst formation to gastrulation. Developmental Cell, 57(2), 152–165. 10.1016/j.devcel.2021.12.022

Rossant, J., & Tam, P. P. L. (2021). Opportunities and challenges with stem cell-based embryo models. Stem Cell Reports, 16, 1031–1038.

Rossi, G., Broguiere, N., Miyamoto, M., Boni, A., Guiet, R., Girgin, M., Kelly, R. G., Kwon, C., & Lutolf, M. P. (2021). Capturing cardiogenesis in gastruloids. Cell Stem Cell, 28(2), 230–240.e6. 10.1016/j.stem.2020.10.013

Rugg-Gunn, P. J., Moris, N., & Tam, P. P. L. (2023). Technical challenges of studying early human development. Development, 150(11). 10.1242/dev.201797

Sanaki-Matsumiya, M., Matsuda, M., Gritti, N., Nakaki, F., Sharpe, J., Trivedi, V., & Ebisuya, M. (2022). Periodic formation of epithelial somites from human pluripotent stem cells. Nature Communications, 13(1), 2325. 10.1038/s41467-022-29967-1

Schnirman, R. E., Kuo, S. J., Kelly, R. C., & Yamaguchi, T. P. (2023). The role of Wnt signaling in the development of the epiblast and axial progenitors. In Wnt Signaling in Development and Disease (Vol. 153, Current Topics in Developmental Biology). Elsevier. 10.1016/bs.ctdb.2023.01.010

Sheng, G. (2021). Defining epithelial-mesenchymal transitions in animal development. Development, 148(8). 10.1242/dev.198036

Simunovic, M., & Brivanlou, A. H. (2017). Embryoids, organoids and gastruloids: New approaches to understanding embryogenesis. Development, 144(6), 976–985. 10.1242/dev.143529

Simunovic, M., Siggia, E. D., & Brivanlou, A. H. (2022). In vitro attachment and symmetry breaking of a human embryo model assembled from primed embryonic stem cells. Cell Stem Cell, 29(6), 962–972.e4. 10.1016/j.stem.2022.05.001

Smirnova, N. P., Ponomartsev, S. V., Liyakat Ali, T. M., et al. (2025). Modular engineering of embryonic-extraembryonic interactions generates advanced gastruloid morphotypes. bioRxiv. 10.1101/2025.11.09.687163

Stapornwongkul, K. S., Hahn, E., Poliński, P., Salamó Palau, L., Arató, K., Yao, L., Williamson, K., Gritti, N., Anlas, K., Osuna Lopez, M., Patil, K. R., Heemskerk, I., Ebisuya, M., & Trivedi, V. (2025). Glycolytic activity instructs germ layer proportions through regulation of Nodal and Wnt signalling. Cell Stem Cell, 32(5), 744–758.e7. 10.1016/j.stem.2025.03.011

Stuart, T., Butler, A., Hoffman, P., Hafemeister, C., Papalexi, E., Mauck, W.M., 3rd, Hao, Y., Stoeckius, M., Smibert, P., and Satija, R. (2019). Comprehensive integration of single-cell data. Cell 177, 1888–1902.e21. 10.1016/j.cell.2019.05.031

Tani, S., Okada, H., Onodera, S., et al. (2023). Stem cell-based modeling and single-cell multiomics reveal gene-regulatory mechanisms underlying human skeletal development. Cell Reports, 42(4), 112276. 10.1016/j.celrep.2023.112276

Turner, D. A., & Martinez-Arias, A. (2024). Three-dimensional stem cell models of mammalian gastrulation. BioEssays, 46, e2400123.

Turner, D. A., Girgin, M., Alonso-Crisostomo, L., Trivedi, V., Baillie-Johnson, P., Glodowski, C. R., Hayward, P. C., Collignon, J., Gustavsen, C., Serup, P., Steventon, B., Lutolf, M. P., & Martinez-Arias, A. (2017). Anteroposterior polarity and elongation in the absence of extra-embryonic tissues and of spatially localised signalling in gastruloids: Mammalian embryonic organoids. Development, 144(21), 3894–3906. 10.1242/dev.150391

van den Brink, S. C., Baillie-Johnson, P., Balayo, T., Hadjantonakis, A.-K., Nowotschin, S., Turner, D. A., & Martinez-Arias, A. (2014). Symmetry breaking, germ layer specification and axial organisation in aggregates of mouse embryonic stem cells. Development, 141(22), 4231–4242. 10.1242/dev.113001

Veenvliet, J. V., Bolondi, A., Kretzmer, H., Haut, L., Scholze-Wittler, M., Schifferl, D., Koch, F., Guignard, L., Kumar, A. S., Pustet, M., Heimann, S., Buschow, R., Wittler, L., Timmermann, B., Meissner, A., & Herrmann, B. G. (2020). Mouse embryonic stem cells self-organize into trunk-like structures with neural tube and somites. Science, 370(6522), eaba4937. 10.1126/science.aba4937

Waaler, J., Leenders, R. G. G., Sowa, S. T., Alam Brinch, S., Lycke, M., Nieczypor, P., Aertssen, S., Murthy, S., Galera-Prat, A., Damen, E., Wegert, A., Nazaré, M., Lehtiö, L., & Krauss, S. (2020). Preclinical lead optimization of a 1,2,4-triazole based tankyrase inhibitor. Journal of Medicinal Chemistry, 63, 6834–6846.

Waaler, J., Mygland, L., Tveita, A., Strand, M. F., Solberg, N. T., Olsen, P. A., Aizenshtadt, A., Fauskanger, M., Lund, K., Brinch, S. A., Lycke, M., Dybing, E., Nygaard, V., Bøe, S. L., Heintz, K., Hovig, E., Hammarström, C., Corthay, A., & Krauss, S. (2020). Tankyrase inhibition sensitizes melanoma to PD-1 immune checkpoint blockade in syngeneic mouse models. Communications Biology, 3(1), 196. 10.1038/s42003-020-0916-2

Wang, W., Xiao, Z., Li, X., Aziz, K. E., Gan, B., Johnson, R. L., & Chen, J. (2015). AMPK modulates Hippo pathway activity to regulate energy homeostasis. Nature Cell Biology, 17(4), 490–499. 10.1038/ncb3113

Williams, M., Burdsal, C., Periasamy, A., Lewandoski, M., & Sutherland, A. (2012). Mouse primitive streak forms in situ by initiation of epithelial to mesenchymal transition without migration of a cell population. Developmental Dynamics, 241(2), 270–283. 10.1002/dvdy.23711

Williams, S. E., Beronja, S., Pasolli, H. A., & Fuchs, E. (2011). Asymmetric cell divisions promote Notch-dependent epidermal differentiation. Nature, 470(7334), 353–358. 10.1038/nature09793

Yamanaka, Y., Hamidi, S., Yoshioka-Kobayashi, K., Munira, S., Sunadome, K., Zhang, Y., Kurokawa, Y., Ericsson, R., Mieda, A., Thompson, J. L., Kerwin, J., Lisgo, S., Yamamoto, T., Moris, N., Martinez-Arias, A., Tsujimura, T., & Alev, C. (2022). Reconstituting human somitogenesis in vitro. Nature, 614(7948), 509–520. 10.1038/s41586-022-05649-2

Zeng, B., Liu, Z., Lu, Y., Zhong, S., Qin, S., Huang, L., Zeng, Y., Li, Z., Dong, H., Shi, Y., Yang, J., Dai, Y., Ma, Q., Sun, L., Bian, L., Han, D., Chen, Y., Qiu, X., Wang, W., … Wang, X. (2023). The single-cell and spatial transcriptional landscape of human gastrulation and early brain development. Cell Stem Cell, 30(6), 851–866.e7. 10.1016/j.stem.2023.04.016

Zheng, Y., Xue, X., Shao, Y., Wang, S., Esfahani, S. N., Li, Z., Muncie, J. M., Lakins, J. N., Weaver, V. M., Gumucio, D. L., & Fu, J. (2019). Controlled modelling of human epiblast and amnion development using stem cells. Nature, 573(7774), 421–425. 10.1038/s41586-019-1535-2

